# Simultaneous Inference of Past Demography and Selection from the Ancestral Recombination Graph under the Beta Coalescent

**DOI:** 10.1101/2022.09.28.508873

**Authors:** Kevin Korfmann, Thibaut Sellinger, Fabian Freund, Matteo Fumagalli, Aurélien Tellier

## Abstract

The reproductive mechanism of a species is a key driver of genome evolution. The standard Wright-Fisher model for the reproduction of individuals in a population assumes that each individual produces a number of offspring negligible compared to the total population size. Yet many species of plants, invertebrates, prokaryotes or fish exhibit neutrally skewed offspring distribution or strong selection events yielding few individuals to produce a number of offspring of up to the same magnitude as the population size. As a result, the genealogy of a sample is characterized by multiple individuals (more than two) coalescing simultaneously to the same common ancestor. The current methods developed to detect such multiple merger events do not account for complex demographic scenarios or recombination, and require large sample sizes. We tackle these limitations by developing two novel and different approaches to infer multiple merger events from sequence data or the ancestral recombination graph (ARG): a sequentially Markovian coalescent (SM*β*C) and a graph neural network (GNN*coal*). We first give proof of the accuracy of our methods to estimate the multiple merger parameter and past demographic history using simulated data under the *β*-coalescent model. Secondly, we show that our approaches can also recover the effect of positive selective sweeps along the genome. Finally, we are able to distinguish skewed offspring distribution from selection while simultaneously inferring the past variation of population size. Our findings stress the aptitude of neural networks to leverage information from the ARG for inference but also the urgent need for more accurate ARG inference approaches.

## Introduction

With the availability of genomes of increasing quality for many species across the tree of life, population genetics models and statistical methods have been developed to recover the past history of a population/species from whole genome sequence data from several individuals [89, 60, 84, 90, 87, 5, 4, 92, 44, 45]. Indeed, the inference of the past demographic history of a species, *i.e.* population expansion, contraction, or bottlenecks, extinction/colonisation, is not only interesting in its own right, but also essential to calibrate genome-wide scans to detect genes under (*e.g.* positive or balancing) selection [92, 46]. A common feature of inference methods that make full use of whole genome sequences is the underlying assumption of a Kingman coalescent process [53] to describe the genealogy distribution of a sample. The Kingman coalescent process and its properties stem from using the traditional forward-in-time Wright-Fisher (WF) model to describe the reproduction mechanism of a population. Besides non-overlapping generations, a key assumption of the neutral WF model is that an individual offspring chooses randomly (*i.e.* uniformly) its parents from the previous generation. More precisely, each chromosome chooses a parental chromosome from the previous generation. Thus, a key parameter is the distribution of the number of offspring that parents can have. In the WF model, due to the binomial sampling, the distribution of offspring number per parent is well approximated by a Poisson distribution with both mean and variance equal to one. This implies that parents will most likely have zero, one, or two offspring individuals, but it is improbable that one parent would have many offspring individuals (*i.e.* on the order of the population size, under the Wright-Fisher haploid model the probability for a parent to have 10 or more offspring is ≈ 10−^8^). The assumption of small variance in offspring distribution between individual parents is realistic for species with low juvenile mortality (so-called type I and II survivorship in ecology, see survivorship curves *e.g.* by [24]), such as mammals.

As genome sequence data become available for a wide variety of species with different biological traits and/or life cycles, the applicability of the Kingman coalescent relying on the WF model can be questioned [91, 2, 3, 71, 47, 68, 94, 65, 33]. Indeed, for some species, such as fish, with high fecundity and high juveniles mortality (type III survivorship, [24]), it is expected that the variance in reproduction between parents can be much larger than under the Poisson distribution [94]. This effect is termed as sweepstake reproduction [38, 2]. Neutral processes such as strong seed banking [12], high fecundity with skewed offspring distribution [38, 28], extremely strong and recurrent bottlenecks [9, 22], and strong selective processes (*i.e.* positive selection) [27, 18, 19, 37, 3] are theoretically shown to deviate from the classic WF model in a way that the genealogies can no longer be described by a Kingman coalescent process. Under such conditions, a new class of processes arise to describe the genealogy distribution, a class where multiple individuals can coalesce and/or multiple distinguished coalescence events can occur simultaneously [80, 67, 26, 79, 73, 14]. Generally, this class of genealogical processes is called the Multiple Merger Coalescent (MMC). MMC models are more biologically appropriate than the Kingman coalescent to study many species of fish [29, 2, 3, 38], invertebrates (insects, crustaceans, etc.), viruses [63], bacteria [65, 69], plants and their pathogens [94]. While we would like to assess which population model best describes the species genealogy, field experiments to quantify the underlying reproduction mechanism of a species can be costly and time consuming at best, or intractable at worst. Therefore, an alternative solution is to use inference methods based on genome data to identify which model best describes the genealogy of a given species/population.

In this study we use the so-called *β*-coalescent, a specific class of MMC models. Unlike under the WF model, under MMC models the ploidy level strongly affects the distribution of genealogies [8]. For simplicity, in this study we focus on haploid organisms. In the polyploid case, where each parent contributes multiple genomes, the SMC formulations of putative intra- and inter-individual coalescence events would need to be carefully modelled, since this effect would lead to smaller coalescence probabilities and a change of the predicted statistical power for demographic inference. It is demonstrated that if the probability of a parent to have *k* or more offspring is proportional to *k^−α^*, where 1 < *α* < 2, then the genealogy can be described by a Λ-coalescent [86]. The latter is a general class of coalescent process describing how and how fast ancestral lineages merge [73, 79]. When using the Beta(2 − *α, α*) distribution as a probability measure for the Λ-coalescent, the transition rates (*i.e.* coalescent rate) can be analytically obtained leading to the *β*-coalescent, a specific MMC model. If *α* tends to 2, then the coalescent process converges to a Kingman coalescent up to a scaling constant as specified in a more detailed way in the documentation of msprime (https://tskit.dev/ msprime/docs/stable/api.html#msprime.BetaCoalescent). The effective population size calculations for the Beta coalescent yield 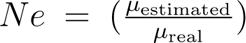 /scaling constant)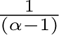, where 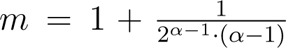, 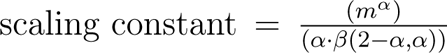 (*β* being the Beta function) and 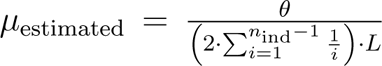 [8, 56, 57, 7, 86] . If *α* tends to one, the model tends to a Bolthausen-Sznitman coalescent process (*i.e.* dominated by strong multiple merger events) [14]. The *β*-coalescent has the property that the observed polarized Site Frequency Spectrum (SFS) of a sample of single nucleotide polymorphisms (SNPs) exhibits a characteristic U-shape with an excess of rare and high frequency variants (compared to the Kingman coalescent) [83]. Current methods to draw inference under MMC models leverage information from the summary statistics extracted from full genome data such as Site Frequency Spectrum (SFS, or derived summary statistics) [57, 37, 78], minor allele frequency [76] or copy number alteration [47]. It is shown that the SFS is robust to the effect of recombination [57, 76] and its shape allows to discriminate between simple demographic models (population expansion or contraction) under the Kingman coalescent and MMC models with constant population size [57, 56, 29]. However, methods relying on genome-wide SFS have two main disadvantages. First, in absence of strong prior knowledge, they can suffer from non-identifiability [44] as several complex neutral demographic and/or selective models under the Kingman or MMC models can generate similar SFS distributions. Second, as they summarize the collection of underlying genealogies, they require high sample sizes (>50) to produce trustworthy results [57, 56, 29], relying on experimental designs which are prohibitive for the study of non-model species. To tackle these limitations, we develop two methods that integrate recombination events along the genome in order to leverage more information from full genome data, thus requiring fewer samples.

In species undergoing sexual reproduction, recombination events break the genealogy of a sample at different position of the genome (*i.e.* the genealogy of a sample varies along the genome), leading to what is called the Ancestral Recombination Graph (ARG) ([41, 8, 102, 59]). Because all the genealogical information is contained in the ARG, in this study we aim at the interpretation of the ARGs to recover model parameters in presence of multiple merger events. With the development of the sequentially Markovian coalescent theory ([64, 62, 101]), it becomes tractable to integrate linkage disequilibrium over chromosomes in inferences based on the Kingman coalescent ([60]). Hence, we first develop an SMC approach based on the *β*-coalescent named the Sequentially Markovian *β* Coalescent (SM*β*C). The *β*-coalescent has the additional property that, under recombination, long range dependency can be generated between coalescent trees along the genome if multiple-merger events happen in a single generation ([8]). In other words, coalescent trees which are located at different places in the genome, and expected to be unlinked from one another ([70]), would show non-zero correlation in their topology and coalescent times. This is because coalescent trees from different genomic regions may all be affected by the same MMC event (merger event of multiple lineages in the past) which then leaves traces in the genome at several loci ([9]). To overcome the theoretically predicted non-Markovian property of the distribution of genealogies along the genome under the *β*-coalescent with recombination ([8]) and the increasing sparsity of genealogies and ancestral nodes with respect to *α* (see Supplementary Figure S18, S19 and S20), we develop a second method based on deep learning (DL) trained from efficient coalescent simulations ([7]). In evolutionary genomics, DL approaches trained by simulations are shown to be powerful inference tools ([89, 55]). Previous work demonstrated that DL approach can help overcome problems mathematically insolvable or computationally intractable in the field of population genetics ([89, 6, 98, 105, 32, 23, 74, 20, 43]). The novelty of our neural network relies on its structure (Graph Neural Network, GNN) and its training algorithm based on the ARG of a sample, or its tree sequence representation ([48, 16, 99]). GNNs are an emerging category of DL algorithm ([17, 103, 21, 108]) that benefit by using irregular domain data (*i.e.* graphs). GNNs are designed for the prediction of node features ([54, 104]), edge features (link prediction) ([107, 85]), or additional properties of entire graphs ([106, 58]). Therefore, GNNs represent a new tool to address the large dimensionality of ARGs, while simultaneously leveraging information from the genealogy (namely topology and age of coalescent events) as a substantial improvement over convolutions of genotype matrices, as currently done in the field ([81]).

We first quantify the bias of previous SMC methods (MSMC and MSMC2 [84, 97]) when performing inference of past population size variation under the *β*-coalescent. We then describe our two methods, SM*β*C and GNN*coal*, and demonstrate their statistical power as well as their respective limitations. From simulated tree-sequence (*i.e.* ARG) and sequence (*i.e.* SNPs) data, we assess the accuracy of both approaches to recover the past variation of population size and the *α* parameter of the Beta-distribution. This parameter indicates how frequent and strong multiple merger events occur (see Supplementary Figure S20). We demonstrate that our approaches can infer the evolutionary mechanism responsible for multiple merger events and distinguish local selection events from genome-wide effects of multiple mergers. We highlight the limits of the Markovian property of SMC to describe data generated under the *β*-coalescent. Finally, we show that both our approaches can model and identify the presence of selection along the genome while simultaneously accounting for non-constant population size, recombination, and skewed offspring distribution. Thus our methods represents a major and necessary leap forward in the field of population genetic inferences.

## Materials and Methods

In our study we first assume the true ARG to be known. Hence, the ARG of the sample is given as input to our methods to estimate recover model parameters of interest (*e.g.* the *α* parameter and/or the past variation of population size). We then show the applicability of our methods by using as input simulated sequence data (*i.e.* SNPs) and/or ARG inferred using ARGweaver [75] from simulated sequence data.

### SMC-based method

In this study, we use different SMC-based algorithms: two previously published, MSMC and MSMC2 [84, 97], and the new SM*β*C. In the latter, the software backbone stems from our previous eSMC [87, 88] whilst the theoretical framework originates from the MSMC algorithm [84] (see Supplementary Text S1). All approaches can either use the ARG or sequence data as input. Providing the ARG as input for MSMC and MSMC2 is enabled by a re-implementation included in the R package eSMC2 previously published in [88]. It is important to mention that there are no theoretical differences in the models whether sequence data or ARG is inputted (see [88] and Supplementary Text S1 for details). The difference is that in one case the hidden states are inferred from sequence data with a forward-backward algorithm, and in the later the sequence of hidden states are directly built from reading the inputted ARG (skipping the forward-backward algorithm). The MSMC2 algorithm focuses on the coalescence time between two haploid samples along the genome. In the event of recombination, there is a break in the current genealogy and the coalescence time consequently takes a new value. A detailed description of the algorithm can be found in [30, 97]. The MSMC algorithm simultaneously analyses multiple sequences (up to 10) and follows the distribution of the first coalescence event in a sample of size *n >* 2 along the sequence based on the Kingman coalescent [53]. A detailed description of MSMC can be found in [84].

Our new approach, SM*β*C, is a theoretical extension of the MSMC algorithm, simultaneously analyzing multiple haploid sequences and focusing on the first coalescence event of a sample size 3 or 4 (this parameter is named *M* throughout the manuscript). We define as *M* the number of lineages simultaneously modeled by either approach. Hence, the SM*β*C follows the distribution of the first coalescence event of a sample size *M* along sequences assuming a *β*-coalescent process. Therefore, our SM*β*C allows for more than two ancestral lineages to join the first coalescence event, or new lineages to join an already existing binary (or triple) coalescent event. Hence, the SM*β*C extends the MSMC theoretical framework by adding hidden states at which more than two lineages coalesce. Currently, the SM*β*C has been derived to analyze for up to 4 sequences simultaneously (due to computational load and mathematical complexity). However the SM*β*C can handle more than M sequences by analyzing all combination of sample size *M* before optimizing the likelihood. The emission matrix is similar to the one of MSMC. As in the MSMC software, the population size is assumed piece-wise constant in time and we discretize time in 40 bins throughout this study. A detailed description of SM*β*C can be found in Supplementary Text S1. To test and validate the theoretical accuracy of our approach, we first study its best case convergence (introduced in [88]) which corresponds to the model’s performance when the true (exact) genealogy is given as input, *i.e.* as if the hidden states are known. Additionally, we also validate the practical accuracy of the SM*β*C on simulated sequence data taking the same input as the MSMC software [84], or using the inferred ARGs by ARGweaver [75]. All SMC approaches used in this manuscript are found in the R package eSMC2 (https://github.com/TPPSellinger/eSMC2).

### GNN*coal* method

Inspired by results obtained from inferences based on tree sequence data [35, 88], we develop a graph neural network (GNN) taking tree sequence data as input. Our GNN is designed to infer population size along with the *α* parameter of the Beta distribution describing the distribution of offspring production. In practice, the ARG is reshaped into a sequence of genealogies (more precisely a sequence of undirected graphs), and then given as input to the GNN (similar to what is described above for the SM*β*C). In our analyses, we fixed the batch size to 500. This value represents the number of coalescence trees being processed before updating parameters of the neural network. As the batch size is fixed to 500, only simulations displaying at least 500 recombination events are considered for the training data sets. If more than 500 recombination events occur along the sequence, the ARG is truncated and the GNN will only take as input the first 500 genealogies and remove the rest. Thanks to the GNN architecture, the algorithm can account for the topology of the genealogy. Hence, the GNN leverages information from coalescence time and branch lengths but also from the topology of the ARG. This operation is known as a graph convolution. By doing so, the GNN is capable of learning from local features of the ARG and extract information from its complex structure. To learn from global genealogy patterns (which SMC-based methods cannot do), an additional pooling strategy is implemented as part of the network [106]. To do so, the ARG is broken into smaller ARGs (*i.e.* subgraphs) during the forward-pass step. To illustrate the GNN strategy, we visualize the compression-like process, from the coalescent trees (1) being processed by GNN*coal* (2,3) to the inferred variable of interest (4, 5) in Figure 1.

**Fig. 1.**
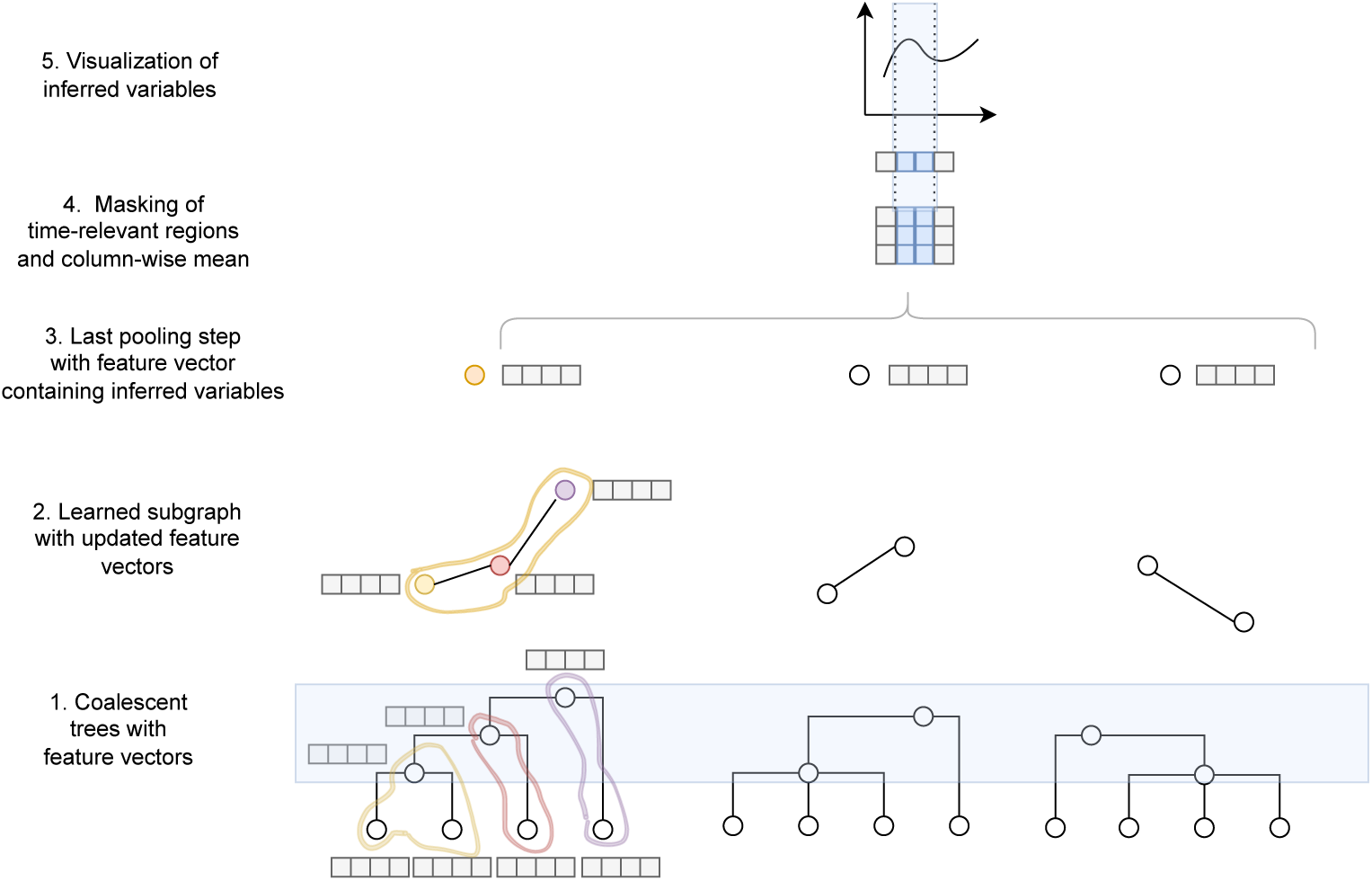
Schematic representation of GNN*coal* processing an ARG. The figure represents the analogues compression of node embeddings (or feature vectors) as in Fig. 1 of [106]. The pooling is hierarchical and applied to each coalescent trees until a single embedding per tree remains, which is fed into a dense neural net to obtain the inferred variable of interest (*i.e.* demographic changes). Each coalescent ancestor or leaf node is initialized by this feature vector (light grey boxes) (1). Sub-graphs are generated by a pooling network with updated feature vectors and a final compression step is performed until ideally one node per graph remains (2-3). Lastly, the column-wise mean is taken after applying a time mask (blue - based on number of coalescent events), so that single feature vector remains (4-5). Detailed description of the graph convolution, feature vector initialization, pooling methodology, coalescent time mask construction, and dataset generation can be found in Supplementary Text S2 or [106].

To infer parameters from our neural network, we need to define an objective function to be optimized. We use a masked root-mean-squared error (RMSE) loss function as objective function which is computed for each inputted ARG (*i.e.* minimizing the average square difference between predicted and true parameter value). In practice, time is discretized (as for the SM*β*C) and time windows are defined. The true *α* value and true demography at 60 predefined time points are given as input to the GNN to compute the loss function. The GNN captures the stochastic complexity arising from the underlying demographic scenario and model parameters. Furthermore, our algorithm naturally defines an appropriate time window to have sufficient observation at each time point. A more detailed description of the GNN*coal* can be found in Supplementary Text S2. The code of the model architecture is implemented in *Pytorch* [72] using the extension *Pytorch Geometric* [31]. The model is available with the simulated training dataset at https://github.com/kevinkorfmann/GNNcoal and https://github.com/kevinkorfmann/GNNcoal-analysis.

### ARGweaver and tsinfer

As the ARG is not known in practice, it needs to be inferred from sequence data. ARG-weaver displays the best performance at recovering the ARG from whole genome polymorphism data at the sample sizes employed in this study (*i.e. « 50*) [75, 15]. Briefly, ARGweaver samples the ARG of *n* chromosomes/scaffolds conditional on the ARG of *n* − 1 chromosomes/scaffolds. To this aim, ARGweaver relies on hidden Markov models while assuming a sequentially Markov coalescent process and a discretization of time, similarly to the SMC-based methods previously described. For a more detail description of the algorithm, we refer the reader to the supplementary material of [75].

For distinguishing between MMC and selection we additionally applied tsinfer to estimate undated genealogical topologies in an effort to build a small training dataset for a model selection study reframed as classification task. Tsinfer has been chosen due to its computational performance and details about the algorithm can be found in the respective supplementary information of [49].

### Simulation of data

#### Validation dataset for both methods

The ARG is given as input to the DL approach and the SM*β*C (see [88]). We use msprime [7] to simulate the ARG of a sample (individuals are assumed to be haploid) under the *β*-coalescent based on [86, 8] or under the Kingman coalescent (under neutrality or selection using msprime *SweepGenicSelection* functionality with start and end frequency of 1/*N_e_* and 0.99, respectively). We simulate 10 sequences of 100 Mbp under five different demographic scenarios: 1) Constant population size; 2) Bottleneck with sudden decrease of the population size by a factor 10 followed by a sudden increase of population by a factor 10; 3) Expansion with sudden increase of the population size by a factor 10, 4) Contraction with sudden decrease of the population size by a factor 10; and 5) “Sawtooth” with successive exponential decreases and increases of population size through time, resulting in continuous population size variation (as shown in [95, 84, 88]). We simulate data under different *α* values (*i.e.* parameters of the *β*-distribution) including values of 1.9 (almost no multiple merger events), 1.7, 1.5, and 1.3 (frequent and strong multiple merger events; Supplementary Figure S20). Mutation and recombination rate (respectively *µ* and *r*) are set to 10^−8^ per generation per bp in order to obtain the best compromise between realistic values and number of SNPs. When specified, some specific scenarios assume recombination and mutation rate set to produce sufficient data or to avoid violation of the finite site hypothesis. All python scripts used to simulate data sets are available at https://github.com/kevinkorfmann/GNNcoal-analysis. Note that the output of msprime suffers from a discontinuity in behaviour when increasing *α* above 1.9 and transitioning from the Beta coalescent to the Kingman coalescent (*α* = 2). The coalescent process converges to a Kingman coalescent up to a scaling constant which we recover in our simulations and estimations (see description in https://tskit.dev/msprime/docs/stable/api.html#msprime.BetaCoalescent).

Additionally, to generate sequence data, we simulate 10 sequences of 10 Mbp under the five different demographic scenarios described above and for the same *α* values. For each scenario, 10 replicates are simulated. In order to obtain sufficient SNPs for inference, we simulate sequence data with mutation and recombination rate (respectively *µ* and *r*) of 10^−8^ per generation per bp when *α* is set to 1.9 and 1.7, 10^−7^ per generation per bp when *α* is set to 1.5, and 10^−6^ per generation per bp when *α* is set to 1.3.

#### Training dataset for the GNN***coal***

In our study we train two GNNs, one to infer past variation of population size through time along with *α*, and one for model selection. The training dataset used for both GNNs is described below.

#### Training dataset for the GNN inferring ***α*** and demography

We generate an extensive number of ARGs to train our GNN. The ARGs are simulated under many demographic scenarios and *α* values. The model parameters are updated in supervised manner. The loss function is calculated for each batch with respect to how much the machine-learning estimates differ from to the true parameters used for simulation. The simulations strategy to recover past demographic history is based on the strategy described and used in [13, 81]. The idea of this approach is to generate a representative set of demographic scenarios over which the network generalizes to consequently infer similar demographic changes after training. More details on the training strategy can be found in Supplementary Text S2.

To improve the simulated demographic history before inference, we introduce a smoothing of the demography allowing to infer continuous variation of population size through time. We do so by interpolating *I* time points cubically, and choosing *w* (set to 60) uniformly spaced new time points of the interpolation in log space. All time points more recent than ten generations in the past are discarded, since inference is too imprecise in the very recent present under our models. An example of this process can be seen in Supplementary Text S2.

#### Training dataset to disentangling coalescent and selection signatures

Beyond parameter inference, deep learning approaches can also be used for clustering. Hence, we train a GNN to disentangle between different scenarios and models. In total, we define eight classes, namely K (S0) (Kingman, no selection), K (WS) (Kingman, weak selection), K (MS) (Kingman, medium selection), K (SS) (Kingman, strong selection) and four different *β*-coalescent classes (1.75 ≤ *α* < 2, 1.5 ≤ *α* < 1.75, 1.25 ≤ *α* < 1.5, 1.01 ≤ *α* < 1.25) without selection. The three different selection regimes are defined as: 0.01 ≤ *Ne* × *s* < 0.1 for SS, 0.001 ≤ *Ne* × *s* < 0.01 for MS, 0.0001 ≤ *Ne* × *s* < 0.001 for WS and *Ne* × *s* = 0 for absence of selection. Demography is kept constant and set to 10^4^ and 10^6^ individuals for Kingman and *β*-coalescent respectively and sequence length is set to 10^5^ bp. The simulation is discarded if it resulted in less than 2,000 obtained trees and is rerun with twice the sequence length until the tree number required is satisfied. This procedure avoids simulating large genome segments of which only a small fraction of trees is used for the given scenario during training and inference. The selection site is introduced in the centre of the respective sequence, so that 249 trees left and 250 right of the middle tree under selection form a training sample, using 500 trees for each sample. One hundred replicates are generated for each training sample. The complete training dataset consists of 4,000 parameter sets: 2,000 for the Kingman cases and 2,000 for the *β*-coalescent cases (90% training dataset and 10% testing dataset). The model itself is trained for 20 epochs (number of time the data is analyzed), and the evaluation performed afterward on 1,000 randomly generated parameter sets, with one replicate per parameter set. Branches of the datasets have been normalized by population size to avoid biases in the dating. Additionally, all tree sequences have been re-inferred with tsinfer to create a separated dataset, which has been used for training and evaluation (see results below). The same architecture used for demography estimation is employed with additional linear layers to reduce the number of output dimensions from 60 to 8. The loss function is set to a Cross-Entropy-Loss for the network to be trainable for categorical labels. Otherwise all architecture and training parameters is the same as described above and detailed in Supplementary Text S2.

## Results

### Inference bias under the wrongly assumed Kingman coalescent

We first study the effect of assuming a Kingman coalescent when the underlying true model is a *β*-coalescent (*i.e.* in presence of multiple merger events) by applying MSMC and MSMC2 to our simulated data. The inference results from MSMC and MSMC2 when the population undergoes a sawtooth demographic scenario are displayed in Figure 2. For *α >* 1.5 the shape of the past demography is fairly well recovered. Decreasing the parameter *α* of the *β*-coalescent (*i.e.* higher probability of multiple merger events occurring) increases the variance of inferences and flattens the demography. Yet, both methods fail to infer the correct population size, due to the scaling discrepancy between the Kingman and *β*-coalescent. While MSMC and MSMC2 assume an underlying Wright-Fisher model as reproduction model, whose genealogy is well approximated by a Kingman coalescent with one unit of coalescent time corresponding to *N* generations, the *β*-coalescent simulation are based on a different reproduction model [86], whose genealogy is given by a *β*-coalescent with a different timescale (see Introduction). Even for *α* close to 2, where the *β*-coalescent resembles the Kingman coalescent, one unit of coalescent time in the *β*-coalescent and one unit in a Wright-Fisher model associated Kingman coalescent still differ by a scaling factor (see Introduction and Methods for details). Hence, we perform the same analysis and correct for the scaling effect after the inference of the MMC versus a Kingman coalescent to better capture the specific effects of assuming binary mergers only. The results are displayed in Figure S1. For *α* > 1.5 the demography is accurately recovered providing we know the true value of *α* to adjust the y-axis (population size) scale. However, for smaller *α* values the observed variance is extremely high and a flattened past variation of population size is observed.

**Fig. 2.**
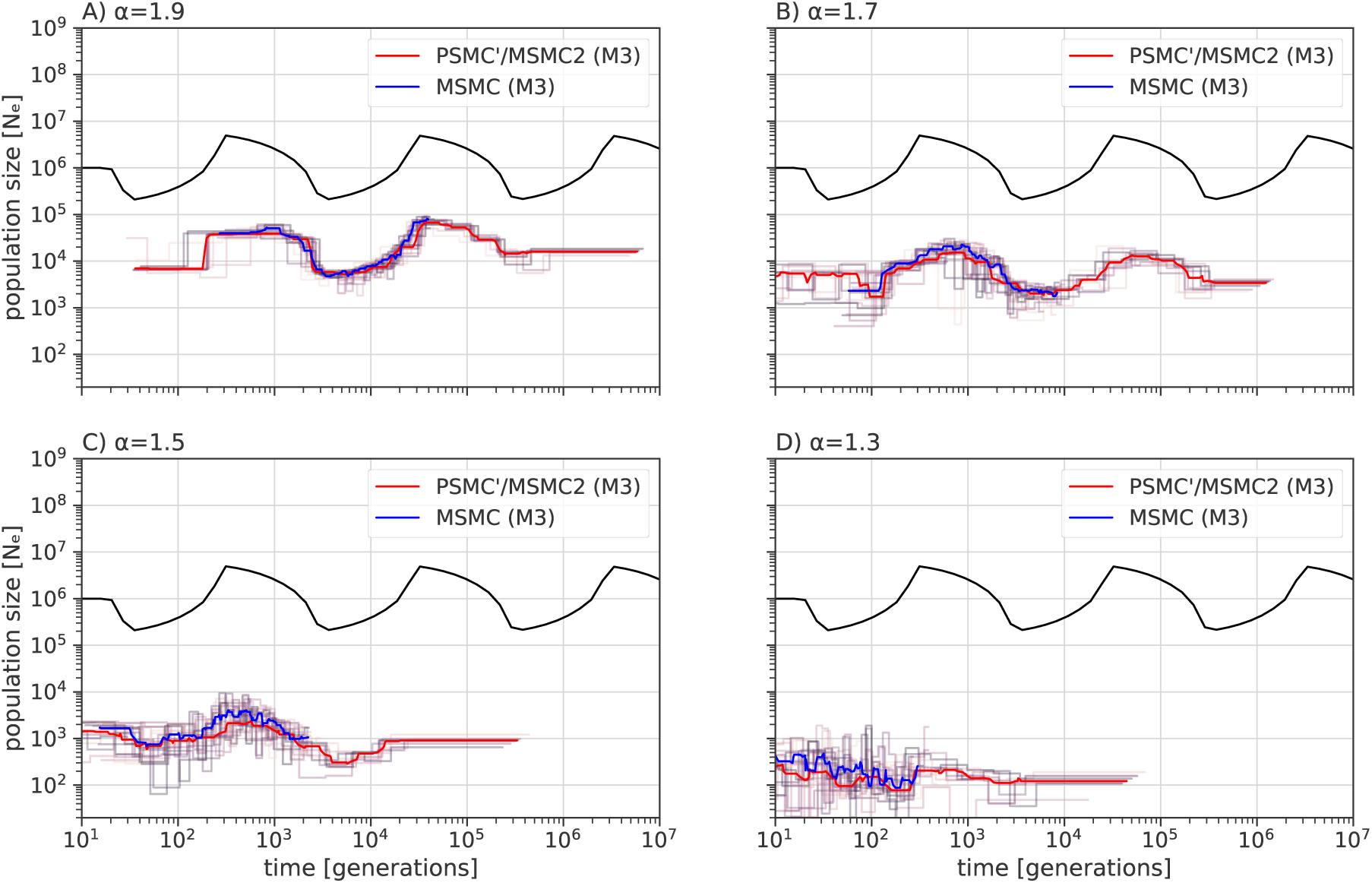
Performance of MSMC and MSMC2 under a *β*-coalescent. Averaged estimated demographic history by MSMC (blue) and MSMC2 (red) based on 10 sequences (mean of random permutations of *M* =3) of 100 Mb with *µ* = *r* = 10^−8^ per generation per bp over ten repetitions (while analyzing simultaneously 3 sequences, noted by M=3). Each repetition result is represented in light red (PSMC’/MSMC2) or in light blue (MSMC). Population undergoes a sawtooth demographic scenario (black) for A) *α* = 1.9, B) *α* = 1.7, C) *α* = 1.5, and D) *α* = 1.3.

### The limit of the Markovian hypothesis

As SMC approaches rely on the hypothesis of Markovian change in genealogy along the genome, we study the effect of *α* on the linkage disequilibrium (LD) of pairs of SNPs (*r*^2^, [77, 66]) in data simulated under the Kingman Coalescent or the *β*-coalescent (with *α* = 1.5 and *α* = 1.3) and constant population size (Figure 3). LD monotonously decreases in average with distance under the Kingman coalescent suggesting the hypothesis of Markovian change in genealogy to be a fair approximation of the genealogical process in that case [100]. Under the *β*-coalescent a similar shape of the distribution is observed but with a higher average amount of LD. We find a higher variance in LD for smaller *α* values. The increased variance results in the occurrence of high spikes of LD along the genome (*e.g.* Figure 3 B). The stochastic increase of linkage along the genome demonstrates that the Markovian hypothesis used to model genealogies along the genome is strongly violated under the *β*-coalescent due to the long range effect of strong multiple merger events [8].

**Fig. 3.**
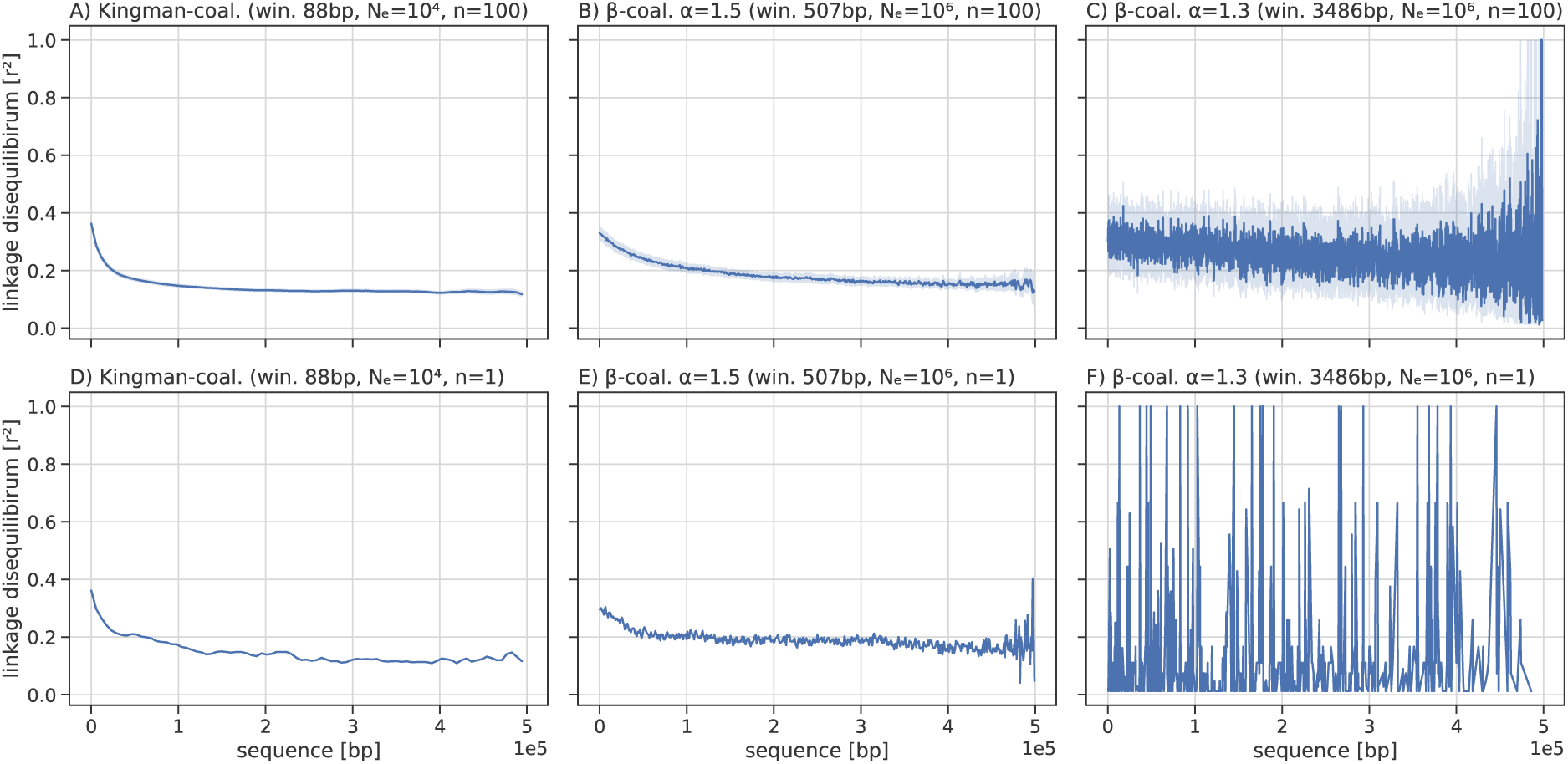
Linkage disequilibrium under a Kingman and *β*-coalescent. Pairwise linkage disequilibrium between SNPs (*r*^2^) under a Kingman and *β*-coalescent with *α* = 1.5 and *α* = 1.3 using 100 sequences of length 0.5 Mb for A) - C) and 1 replicate in D) - F). The population size is constant at *N* = 10^4^ for the Kingman model and *N* = 10^6^ for the *β*-coalescent, with *µ* = 1 × 10 −^7^ and *r* = 1 × 10−^8^ per generation per bp. For each LD analysis, the linkage disequilibrium is calculated by averaging it over automatically-selected window sizes, such that on average at least two mutations are in each window for A) to F), respectively.

We further investigate the effect of multiple merger events on LD. To this aim, we first assume an SMC framework (*e.g.* MSMC2 or eSMC) to predict the transition matrix (*i.e.* matrix containing the probabilities for the coalescent time to change to another value between two positions of the genome) and investigate the absolute difference between the observed transition events. Under the Kingman coalescent, the distribution of coalescent times between two positions in a sample of size two (*n* = 2) is well spread across hidden states in Figure S2 (*i.e.* absence of structured difference between observed and predicted transition events). However, under the *β*-coalescent (with *α* = 1.3) we observe significant differences between observed and predicted transition events at times points where multiple merger events occur (Figure S3). More precisely we observed transitions at specific time points (corresponding to multiple merger events) occurring much more frequently than what is predicted by the model (dark blue lines). This plot thus shows that multiple merger events do not affect the genealogy at every time point and that multiple merger events are over represented in the distribution of transitions events due to the long range effects of MMC events (*i.e.* many positions of the genome contain the same information). This means that one multiple merger coalescent events can affect all positions in the genomes (explaining the spikes in the LD distribution). In contrast, under the Kingman coalescent with recombination, the probability for a coalescent event to affect the whole genome is negligible.

This plot thus unveils the discrepancy between the expectation from the SMC (*i.e.* approximating the distribution of genealogies along the genome by a Markov chain) and the actual effect of multiple merger events on the genealogy distribution along the genome. This discrepancy does not stem from the simulator, because it correctly generates ARG under the *β*-coalescent model [8, 7], but from the limits of the SMC approximation to model events with long range effects on the ARG (Figure S3).

### Inferring ***α*** and past demography on ARG

To test if our two approaches (GNN*coal* and SM*β*C) can recover the past variation of population size and the *α* parameter, we run both methods on simulated tree sequences under different *α* values and demographic scenarios. Figure 4 displays results for data simulated under a sawtooth past demography and for *α* ranging from 1.9, 1.7, 1.5 to 1.3. In all cases, the GNN*coal* approach exhibits low variance to infer the variation of population size and high accuracy from 1.9 to 1.5 with a noticeable drop in accuracy for 1.3 attributable to the ever increasing sparsity due to decreasing *α* generating stronger *β*-coalescent events. For high *α* values (>1.5), the shape of population size variation is well recovered by SM*β*C (4). However, for smaller values, the observed high variance demonstrates the limits of SMC inferences.

**Fig. 4.**
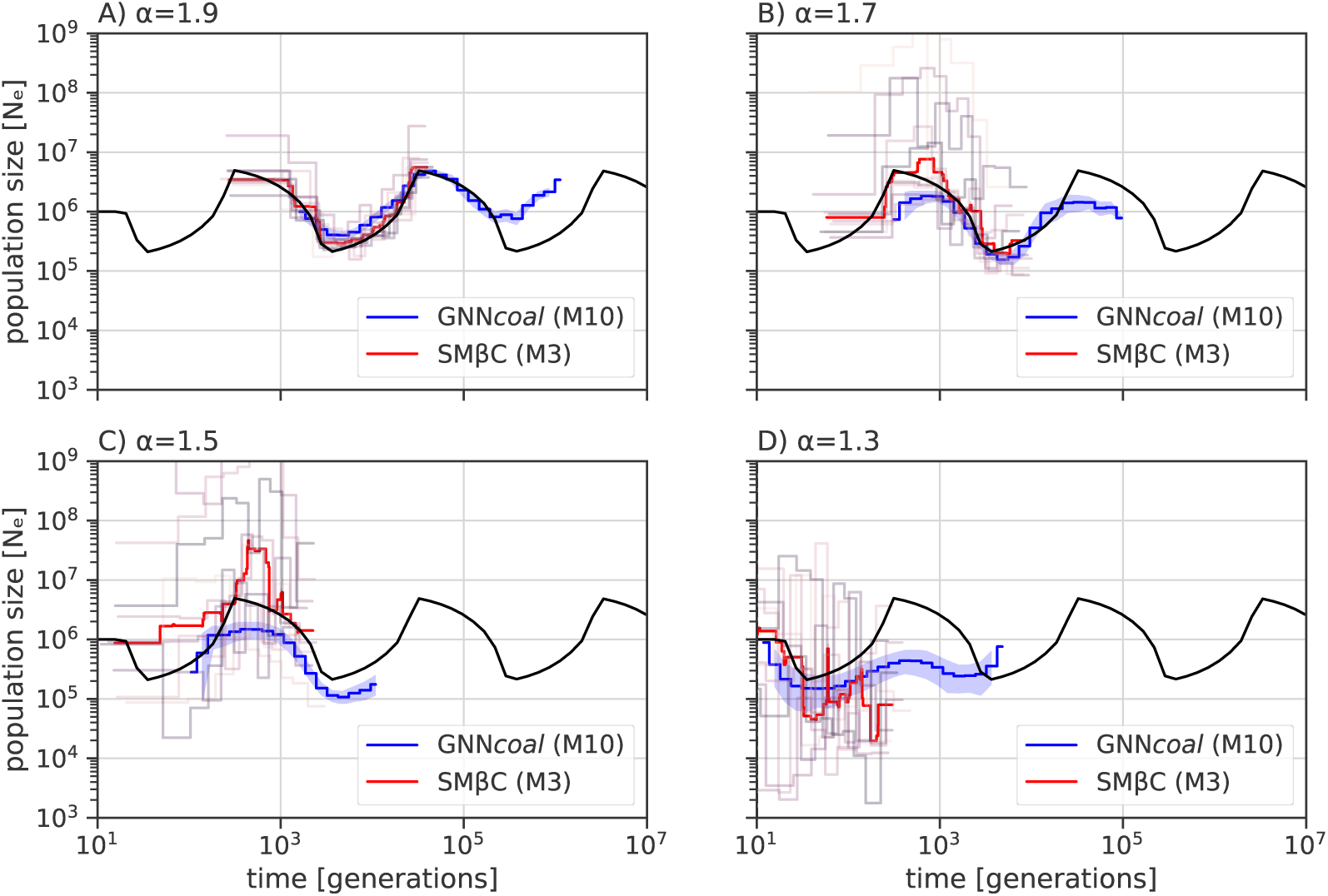
Best-case convergence estimations of SM*β*C and GNN*coal* under a *β*-coalescent. Estimations of past demographic history by SM*β*C in red (median) and by GNN*coal* in blue (mean and 95% confidence interval, CI95; while analyzing simultaneously *M* =3 or *M* =10 sequences; individual replicates of SM*β*C shown as light lines) when population undergoes a sawtooth demographic scenario (black) under A) *α* = 1.9, B) *α* = 1.7, C) *α* = 1.5 and D) *α* = 1.3. SM*β*C runs on 10 sequences and 100 Mb, GNN*coal* runs on 10 sequences and 500 trees, and *µ* = *r* = 10^−8^ per generation per bp.

On average, both approaches seem to recover fairly well the true *α* value (Figure 5 and Table S1). In particular, GNN*coal* displays high accuracy and lower standard deviation. We note that the variance in the estimation of *α* increases with diminishing *α* value. Moreover, increasing the number of simultaneously analyzed sequences by SM*β*C does not seem to improve the inferred *α* value (Table S1). These conclusions are also valid for the results in Figure S4-S7 and Table S1 based on inference under four additional demographic scenarios: constant population size, bottleneck, sudden increase and sudden decrease of population size.

**Fig. 5.**
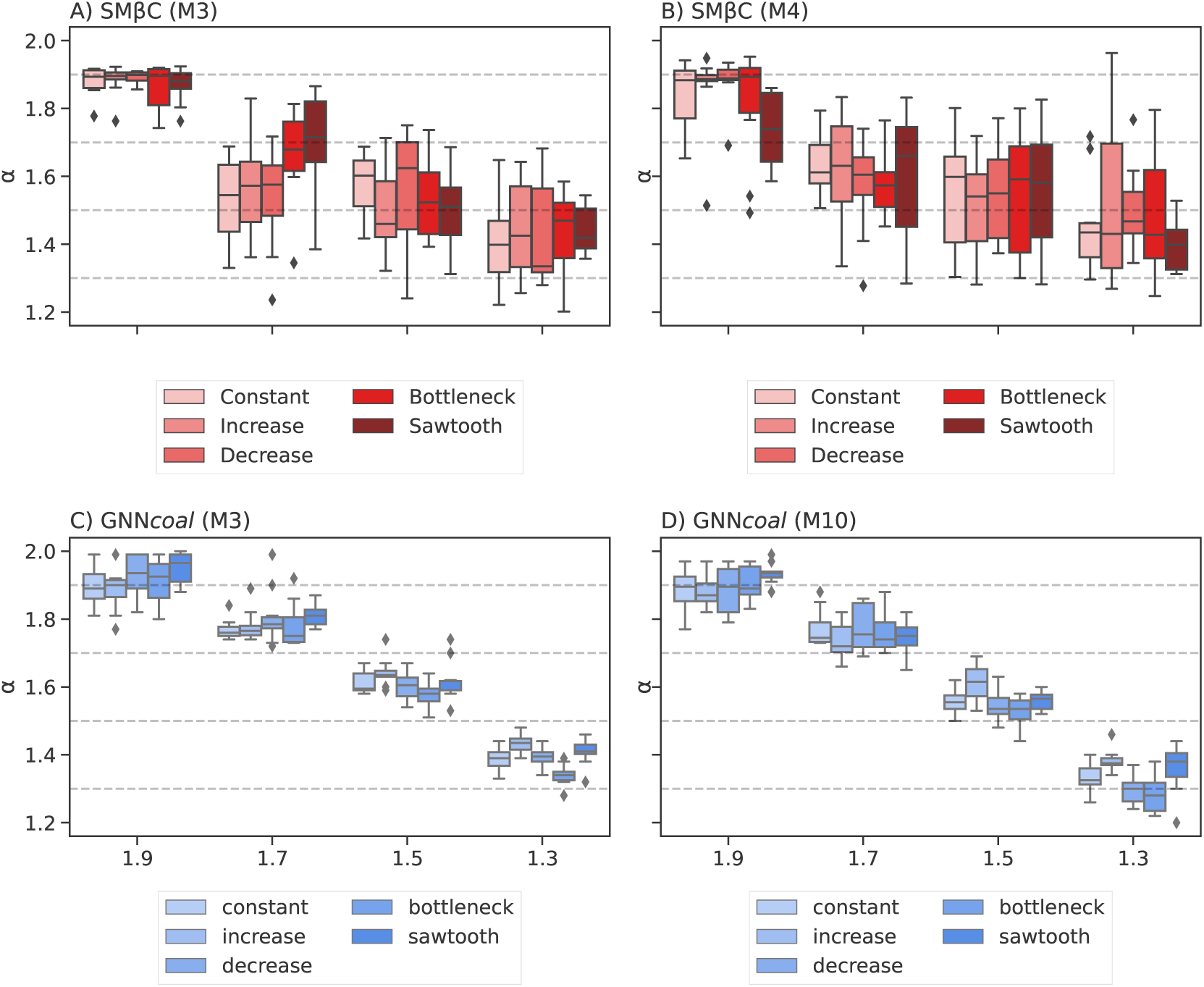
Estimated *α* values by SM*β*C and GNN*coal*. Estimated values of *α* by SM*β*C and GNN*coal* over ten repetitions using 10 sequences of 100 Mb with *µ* = *r* = 10^−8^ per generation per bp under a *β*-coalescent process (with different *α* parameter). The analysis are run on five different demographic scenarios (Constant population size, Bottleneck, Sudden increase, Sudden decrease and a Sawtooth demography) using a sample size *n* = 3 for A) and C), *n* = 4 for B), and *n* = 10 for D). Grey dashed lines indicate the true *α* values. For exact values and standard deviations of the respective experiment see Supplementary Table S1.

When *α* diminishes, the effective population size decreases and the number of recombination events plummets for small values of *α <* 1.5. To demonstrate the theoretical convergence of SM*β*C to the correct values, we run SM*β*C on data simulated with mutation and recombination rate fifty times higher under similar scenarios as in Figure 4. This operation increases the amount of data in the form of SNPs and number of independent coalescent trees by recombination. Since branch lengths (in generations) are on average smaller in the presence of multiple merger when compared to a Kingman coalescent, we choose to increase the rates as opposed to increasing the genome lengths, which does not affect the branch lengths (but increases the number of genealogies). Results of SM*β*C for *α* values of 1.7, 1.5 and 1.3 are displayed on Table S2. Overall our results show that SM*β*C can recover *α* with higher accuracy when more data is available. To be more precise when *M* = 3 (*M* being the number of simultaneously haploid sequence analyzed), the overall average inferred *α* values improve from 1.6, 1.53 and 1.42 (Table S1) to 1.64, 1.49 and 1.36 (for data simulated respectively under *α* = 1.7,*α* = 1.5 and *α* = 1.3). Yet when *M* = 4 a gain in accuracy is only observed for *α* = 1.5 and *α* = 1.3. Indeed, the overall average inferred *α* values changed from 1.60, 1.54 and 1.47 (Table S1) to 1.58, 1.47 and 1.39 (for data simulated respectively under *α* = 1.7, *α* = 1.5 and *α* = 1.3).

Although 10 sequences are given to SM*β*C in the previous analyses, the method can only analyze three or four simultaneously. On the other hand, GNN*coal* can simultaneously analyze 10 sequences, that is the whole simulated ARG. As we observe that GNN*coal* has a higher performance than SM*β*C, we wish to test whether the GNN*coal* better leverages information from the ARG or benefits from simultaneously analyzing a larger sample size. Thus, we run GNN*coal* on the same dataset, but downsampling the coalescent trees to a sample size three. Results for sample size ten are displayed in Figure S4 to S7 and downsampled results with sample size three (*M* =3) of GNN*coal*, which appear to be similar, are displayed in Figure S8, demonstrating that the GNNs can better leverage information from the ARG in presence of multiple merger events.

Additionally, we test if both approaches can recover a Kingman coalescent from the ARG when data are simulated under the Kingman coalescent, namely both approach should recover *α* = 2. To do so, we simulate the same five demographic scenarios as above under a Kingman coalescent and infer the *α* parameter along with the past variation of population size. Estimations of *α* values are provided in Table 1 and are systematically higher than 1.85, suggesting mostly binary mergers. The associated inferred demographies are shown in Figures S9-S13. Both approaches correctly infer the past demographic shape up to the scaling discrepancy between the Beta and the Kingman coalescent (as previously described). Furthermore, we notice that the scaling effect only affects the y-axis for the SM*β*C but affect both axes for GNN*coal*.

**Table 1:** Average estimated values of *α* by SM*β*C and GNN*coal* over ten repetitions under the Kingman coalescent using 10 haploid sequences of 10 Mb and *µ* = *r* = 10^−8^ per generation per bp. The standard deviation is indicated in brackets.

| scenario | True $\alpha$ | $\alpha$:SMβC,M=3 | $\alpha$:SMβC,M=4 | $\alpha$: GNN, M=3 | $\alpha$: GNN, M=10 |
| --- | --- | --- | --- | --- | --- |
| Constant | 2 | 1.97 (0.005) | 1.97 (0.008) | 1.99 (0.002) | 1.99 (0.003) |
| Sawtooth | 2 | 1.94 (0.017) | 1.87 (0.019) | 1.99 (0.002) | 1.99 (0.003) |
| Bottleneck | 2 | 1.97 (0.01) | 1.97 (0.009) | 1.99 (0.003) | 1.99 (0.004) |
| Decrease | 2 | 1.97 (0.007) | 1.97 (0.008) | 1.99 (0.003) | 1.99 (0.004) |
| Increase | 2 | 1.97 (0.007) | 1.97 (0.008) | 1.99 (0.004) | 1.99 (0.002) |

As GNN*coal* was not trained on data simulated under the Kingman coalescent (especially with such high population size), some events fall beyond the scope of the GNN due to the scaling discrepancy between the Beta and Kingman coalescence. Hence, we run GNN*coal* on data simulated under the Kingman coalescent but with smaller population size (scaled down by a factor 100) to assure that all events fall within the scope of the GNN. Values of *α* inferred by the GNN*coal* and the SM*β*C under the five demographic scenarios are available in Table S3. The associated inference of population size are plotted in Figure S9-S12. Both approaches recover high *α* values (*i.e.*>1.85) suggesting a genealogy with almost exclusively binary mergers. In addition, both approaches accurately recover the shape of the past variation of population size up to a scaling constant but only on the population size y-axis.

### Inferring ***α*** and past demography from simulated sequence data

We first investigate results for both GNN*coal* and SM*β*C with the objective of evaluating the performance on ARG reconstructed from sequence data using ARGweaver [75] as ARGweaver is currently being considered the best performing approach to infer ARG for sample size smaller than 20 [15]. Demographic inference results by both approaches are displayed in Figure S14, and *α* inference results in Table S4. GNN*coal* does not recover the shape of the demographic history from the inferred ARGs and largely overestimates *α*. In contrast, SM*β*C produces better inferences of *α* when giving the inferred ARG as input when compared to the GNN. SM*β*C recovers the shape of the past variation of population size for *α >* 1.3 but displays extremely high variance for *α* = 1.3. We then evaluate SM*β*C on simulated sequence data to compare the necessity of reconstructing the ARG for the SMC method and found that *α* is typically well recovered (Table 2) and that results are similar to what obtained when the true ARG is given. Furthermore, the shape of the past variation of population size is well inferred under the sawtooth demographic scenario for *α >* 1.3 (Figure S15). In the other four scenarios, the shape of the demography is recovered in recent times but population sizes are underestimated in the past (Figure S16). Finally, as found above from inputted ARGs, the variance in estimates of population sizes generally increases with diminishing *α*.

**Table 2:** Average estimated *α* values by SM*β*C on simulated sequence data over ten repetitions using 10 sequences of 10 Mb with recombination and mutation rate set to 1 × 10^−8^ for *α* 1.9 and 1.7, 1 × 10^−7^ for *α* 1.5 and 1 × 10^−6^ for *α* 1.3 per generation per bp under a Beta coalescent process. The analysis are run on five different demographic scenarios (Constant population size, Bottleneck, Sudden increase, Sudden decrease and a Sawtooth demography).

| scenario | True $\alpha$ | $\alpha^*$:SMβC,M=3 |
| --- | --- | --- |
| Constant | 1.9 | 1.86 (0.16) |
| Bottleneck | 1.9 | 1.89 (0.09) |
| Increase | 1.9 | 1.93 (0.07) |
| Decrease | 1.9 | 1.96 (0.04) |
| Sawtooth | 1.9 | 1.76 (0.17) |
| Constant | 1.7 | 1.82 (0.10) |
| Bottleneck | 1.7 | 1.64 (0.23) |
| Increase | 1.7 | 1.82 (0.10) |
| Decrease | 1.7 | 1.89 (0.13) |
| Sawtooth | 1.7 | 1.71 (0.27) |
| Constant | 1.5 | 1.52 (0.30) |
| Bottleneck | 1.5 | 1.64 (0.33) |
| Increase | 1.5 | 1.57 (0.24) |
| Decrease | 1.5 | 1.60 (0.18) |
| Sawtooth | 1.5 | 1.66 (0.14) |
| Constant | 1.3 | 1.31 (0.20) |
| Bottleneck | 1.3 | 1.2 (0.17) |
| Increase | 1.3 | 1.24 (0.13) |
| Decrease | 1.3 | 1.57 (0.11) |
| Sawtooth | 1.3 | 1.37 (0.16) |

### Inferring MMC and accounting for selection

As specific reproductive mechanisms and selection can lead to the occurrence of multiple merger-like events, we train our neural network on data simulated under the *β*-coalescent, and under the Kingman coalescent in presence or absence of selection to assess our methods capacity to distinguish between them. We then use the trained GNN*coal* to determine if multiple merger events originate from skewed offspring distribution or positive selection, or if the data follows a neutral Kingman coalescent process. The classification results are displayed in Figure 6 in the form of confusion matrices, where the percentage of times the GNN*coal* correctly assigns the true model shown on the diagonal evaluated on a test dataset of 1,000 ARGs. We tested three scenarios A) training and evaluating on known exact ARGs, B) training on exact ARGs but evaluating on inferred ARGs, and, lastly C) training and evaluating on inferred ARGs. The results indicate the necessity of integrating inference errors or instances of branch unresolvability into the training process. The network is able of distinguishing between signals of multiple merger, which translate to an estimate of *α*, from simple ARG-estimation uncertainties. The overall confusion between neighboring classes may be attributed to the comparably small size of training data (4,000 simulations), which enabled to build a training dataset comprised of inferred trees within few hours. To summarize our approach can accurately distinguish between Kingman and *β*-coalescent, but uncertainty needs to be part of the training procedure.

**Fig. 6.**
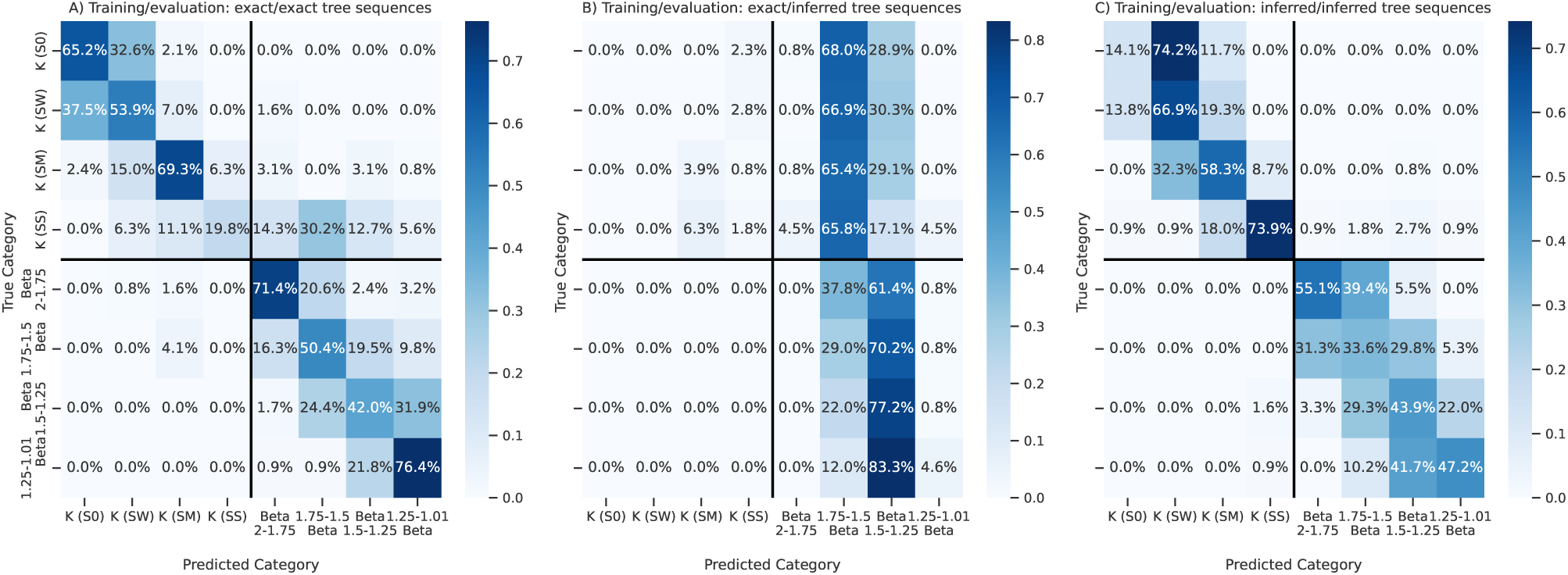
Confusion matrix for Kingman and *β*-coalescent classification model under varying selection coefficients. Evaluation of classification accuracy for Kingman (K) and *β*-coalescent (B) for no selection (S0), weak selection (SW), medium selection (SM) and strong selection (SS) using a 1,000 repetition validation dataset (and small 4000 proof-of-concept repetition training set). Population size was kept constant at *N* = 10^4^ individuals for the Kingman scenario and at *N* = 10^6^ for the *β*-coalescent, using a sample size *n* = 10 and *r* = 10^−8^ per bp per generation. Branch length are normalized by the respective population size. Classification model has been trained and evaluated either on exact or inferred tree sequences (tsinfer without dating) as indicated in the subfigure titles of A), B) and C).

Since strong selection can lead to multiple merge coalescent or rapid and successive coalescent events (as the beneficial alleles spreads very quickly in the population) [27, 11, 78], we investigate if our approaches can model and recover the effect of selection. Therefore, we infer *α* along the genome (to model the local effect of selection on the genome) with both approaches from true genealogies simulated with strong positive selection or neutrality under a Kingman coalescent with population size being constant through time. SM*β*C infers *α* on windows of 10kbp along the genome, and GNN*coal* infers *α* every 20 trees along the genome. Results for GNN*coal* and SM*β*C are displayed in Figure 7. The SM*β*C approach recovers smaller *α* value around the locus under strong selection (while GNN*coal* displays higher variance). However under neutrality or weak selection, inferred *α* values remain high (>1.6) along the genome.

**Fig. 7.**
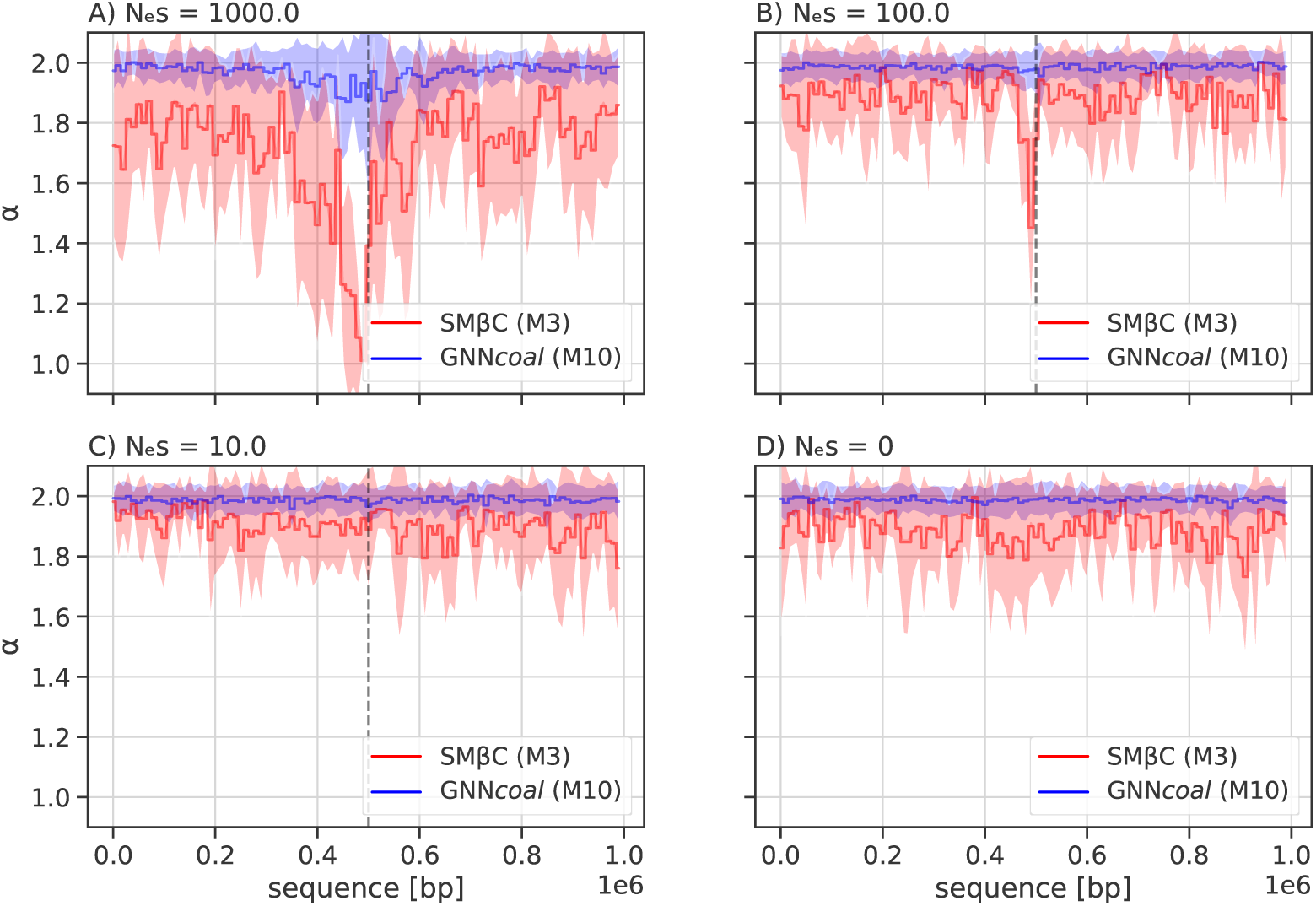
Averaged estimations by GNN*coal* and SM*β*C under selection. Estimations of *α* along the genome by the GNN*coal* approach and the SM*β*C when population undergoes as strong positive selective sweep event (at position 0.5 Mb) under different strengths of selection: A) *s* = 0.01, B)*s* = 0.001, C) *s* = 0.0001, and D) *s* = 0 meaning neutrality (mean and standard deviation for both methods). The population size is constant and set to *N* = 10^5^ with *µ* = *r* = 10^−8^ per generation per bp. We hence have in A) *N_e_* × *s* = 1000,B) *N_e_* × *s* = 100, C) *N_e_* × *s* = 10 and D) *N_e_* × *s* = 0. SM*β*C uses 20 sequences of 1Mb (red) and GNN*coal* uses 10 sequences through down-sampling the sample nodes (blue)

Similarly, we run both approaches on genealogies simulated under the *β*-coalescent (assuming neutrality) and we infer the *α* value along the genome. Inferred *α* values by both approaches are plotted in Figure S17. GNN*coal* is able to recover the *α* value along the genome with moderate overestimation due to tree sparsity. On the contrary, SM*β*C systematically underestimates *α* values. Nevertheless, unlike in presence of positive selection at a given locus, the inferred *α* values are found in all cases to be fairly constant along the genome.

We finally simulate data under a Kingman coalescent (true genealogies) with a strong selective sweep or under neutrality conditioned on a sawtooth demographic scenario to test our methods’ simultaneous inference capabilities. Under neutrality, our both approaches recover, as expected, high *α* values along the genome and can accurately recover the past variation of population size (only up to a scaling constant for GNN*coal*, since it was trained on the *β*-coalescent only) (Figure 8). Similarly, when the simulated data contains strong selection, a small *α* value is recovered at the locus under selection and the past variation of population size is accurately recovered, albeit with a small underestimation of population size in recent times for SM*β*C (Figure 8).

**Fig. 8.**
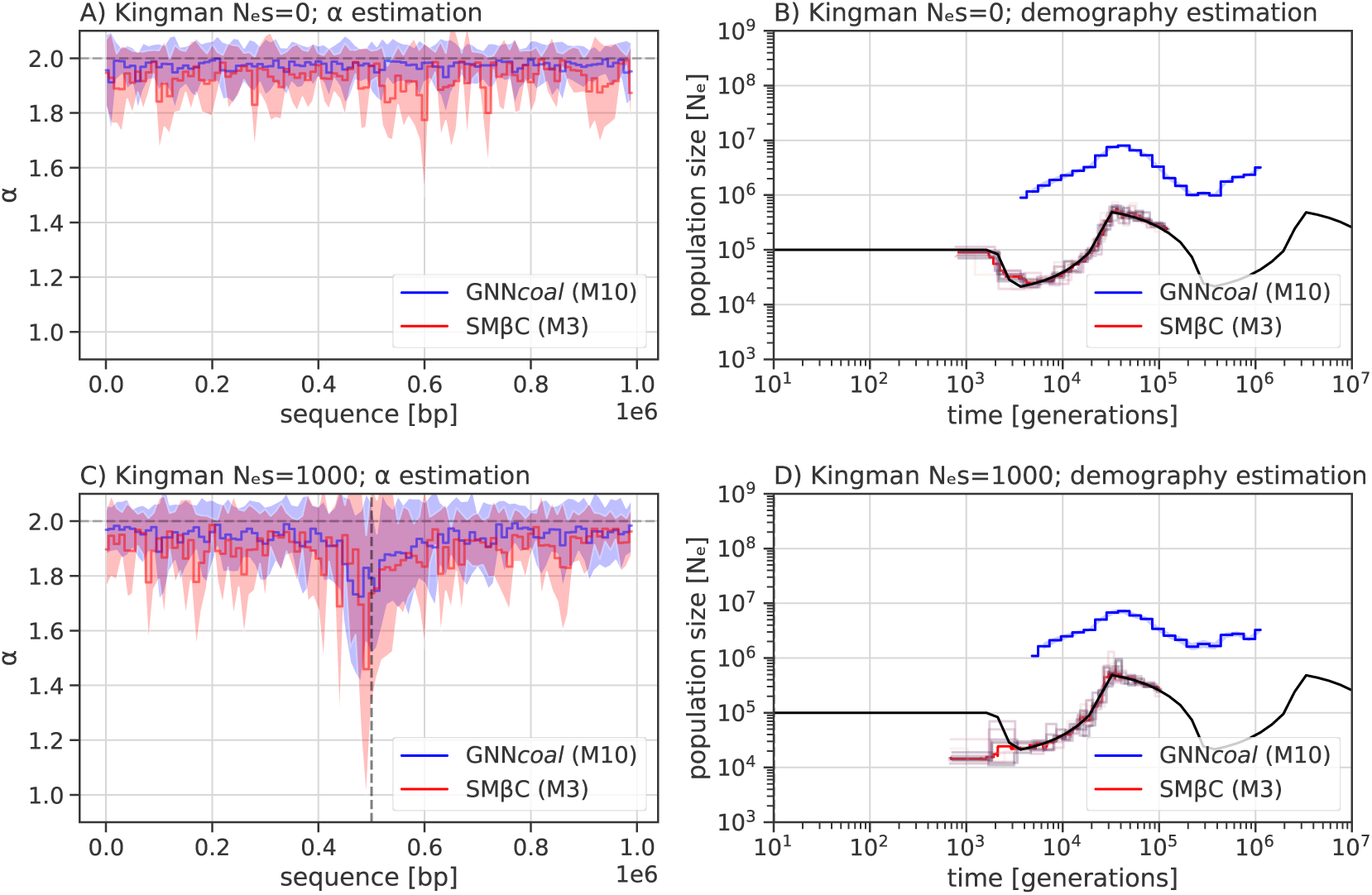
Simultaneous estimations of *α* along the sequence under demographic change by GNN *coal* and SM*β*C. Simultaneous estimation of *α* along the genome under a partial sawtooth scenario: A) and B) in the absence of selection (mean and standard deviation for both methods), and C) and D) presence of selection with *N_e_S* = 1, 000 (mean and CI95 for GNN*coal* and median for SM*β*C). SM*β*C uses 20 sequences of 1Mb (red) and GNN*coal* uses 10 sequences through down-sampling the sample nodes (blue), and *µ* = *r* = 10^8^ per generation per bp.

## Discussion

With the rise in popularity of SMC approaches for demographic inferences [60], most current methods leverage information from whole genome sequences by simultaneously reconstructing a portion of the ARG to infer past demographic history [60, 84, 95, 96], migration rates [52, 97], variation in recombination and mutation along the genome [5, 4], as well as ecological life history traits such as selfing or seed banking [87, 93]. However, other previous studies proposed to uncouple both steps, namely by first reconstructing the ARG and by then inferring parameters from its distribution [88, 35, 75]. Indeed, recent efforts have been made to improve approaches to recover the ARG [90, 50, 40, 75, 61, 15], as well as its interpretation [34, 88]. Our results on data simulated under the *β*-coalescent clearly show the strong effect of multiple merger events on the topology and branch length of the ARG. We find that the more multiple merger events occur, the more information concerning the past demography is lost. Both GNN*coal* and SM*β*C, whether given sequence data, the true or inferred ARG, can recover the *α* parameter and the variation of past population size for *α* values high enough (*i.e. α* ≥ 1.5). However, for lower values of *α*, a larger amount of data is necessary for any inference, specifically in the form of a high effective population size (correspondingly adequate mutation and recombination rates) and sufficient sequence length, which becomes nearly impossible when *α* tends to one. Both approaches can also recover the Kingman coalescent (*i.e. α*>1.8). We find that GNN*coal* outperforms SM*β*C in almost all cases when given the true ARG, and we demonstrate that GNN*coal* can be used to disentangle between *β*-coalescent and Kingman models with selection.

Overall, our results provide a substantial improvement in the development of inference methods for models with multiple merger events, a key step to understand the underlying reproduction mechanism of a species. While still inferring population sizes of the correct order of magnitude, SM*β*C is outperformed by GNN*coal* when given true ARGs as input. As ARG inference method improve, GNN models will offer a promising alternative to current SMC methods. As we directly compare our theoretical SMC to the GNN based on the same input data (coalescent trees), we are ideally placed to dissect the mechanisms underlying the power of the GNN*coal* method. We identify four main reasons for the difference in accuracy between the two methods developed. First, the SM*β*C approach suffers from the limit of the sequential Markovian coalescent hypothesis along the genome when dealing with strong multiple merger events [8, 22]. Second, most current SMC approaches, except XSMC [51], rely on a discretization of the coalescent times into hidden states, meaning that simultaneous mergers of three lineages may not be easily distinguished from two consecutive binary mergers occurring over a short period. Third, the SM*β*C relies on a complex hidden Markov model and due to computational and mathematical tractability, it cannot leverage information on a whole genealogy. In fact, as MSMC, SM*β*C only focuses on the first coalescent event, and therefore cannot simultaneously analyze large sample size. Furthermore, the SM*β*C approach leverages information from the distribution of genealogies along the genome. Whilst, in the near absence of recombination events, both approaches cannot utilize any information from the genealogy itself, GNN*coal* can overcome this limit by increasing the sample size. Fourth, the SM*β*C is based on a coalescent model where *α* is constant in time. Yet multiple merger events do not appear regularly across the genealogical timescale, but occur at few random time points. Hence, the SMC approach suffers from a strong identifiability problem between the variation of population size and the *α* parameter (for low *α* values). For instance, if during one hidden state one strong multiple merger event occurs, multiple merger events are seldom observed and SM*β*C may rather assume a small population size at this time point (hidden state). This may explain the high variance of inferred population sizes under the *β*-coalescent.

By contrast, GNN*coal* makes use of the whole ARG, and can easily scale to larger sample sizes (over 10), although it recovers *α* with high accuracy with sample size *M* = 3 only. Our interpretation is that GNN*coal* is able of simultaneously leveraging information from topology and the age of coalescent events (nodes) across several genealogies (here 500). GNN*coal* ultimately leverages information from observing recurrent occurrences of the same multiple merger events at different locations on the genome, while being aware of true multiple merger events from rapid successive binary mergers. We believe that our results pave the way towards the interpretability of GNN and deep learning methods applied to population genetics. For further theoretical insights into recent descriptions of multiple merger we would like to point the reader towards [25].

When applying both approaches to simulated sequence data (and not to true ARGs), both approaches behave differently. GNN*coal* is not capable to accurately infer model parameters, *i.e.* past variation of population size or *α*. In contrast, SM*β*C performed better than GNN*coal* when dealing with sequence data (and not true ARG). SM*β*C is capable of recovering *α* and the shape of the demographic scenario in recent times irrespective of whether sequence data or ARG inferred by ARGweaver is given as input. This is most likely because the statistic used by SM*β*C (*i.e.* first coalescent event in discrete time) is coarser than the statistic used by GNN*coal* (*i.e.* the exact ARG). We therefore speculate that the theoretical framework of the SM*β*C, although being in theory less accurate than GNN*coal*, is more robust and suited for application to sequence data. More specifically, the issue being faced by the GNN*coal* is known as out-of-distribution inference [42], which requires the network to generalize over an untrained data distribution. This issue happens because GNN*coal* is not trained using ARG inferred by ARGweaver. Building a training data set for GNN*coal* to overcome this issue is currently impractical due to the inference speed of ARGweaver. However, future work will aim at increasing robustness of GNN inferences, for instance by adding uncertainty or multiple models during the training process. Improving the performance of GNN*coal* on sequence data requires more efficient and accurate ARG inference methods, such as to incorporate inferred (non-exact) genealogies into the training, thereby accounting for inference errors and for the evaluation of the algorithm on a broader spectrum of common population genetic research questions. The former observation is important to avoid bias from potential hypothesis violations of the chosen ARG inference approach.

Past demographic history, reproductive mechanisms, and natural selection are among the major forces driving genome evolution [44]. Hence, in the second part of this manuscript we focus on integrating selection in both approaches. Currently, no method (especially if relying only on SFS information) can account for the presence of selection, linkage disequilibrium, non-constant population size and multiple merger events [44] although recent theoretical framework might render this possible in the future [1]. As a first step to fill this gap, we demonstrate that GNN*coal* can be used for model selection to reduce the number of hypotheses to test. Determining which evolutionary forces are driving the genome evolution is key, as only under the appropriate neutral population model results of past demography and selection scans can be correctly interpreted [44, 46]. The high accuracy of GNN*coal* in model selection is promising, especially as other methods based on the SFS alone [57, 47] have limits in presence of complex demographic scenarios. GNN can possibly overcome these limits, as it is easier to scale the GNN to estimate more parameters. We follow a thread of previous work [78, 39, 11], by integrating and recovering selection, multiple merger and population size variation by simply allowing each fixed region in the genome to have its own *α* parameter. In presence of strong selection, we find lower *α* value around the selected loci and high *α* value in neutral neighbouring regions. Hence, our results point out that strong selection can indeed be modeled as a local multiple merger event (see [27, 11, 78]). In presence of weak selection, no effect on the estimated *α* value is observed, demonstrating that weak selection can be modeled by a binary merger and has only a local effect on the branch length by shortening it. In theory, both approaches should be able to infer the global *α* parameter linked to the reproductive mechanism, as well as the local *α* parameter resulting from selection jointly with the variation of population size. However, the absence of a simulator capable of simulating data with selection and non-constant population size under a *β*-coalescent model prevents us from delivering such proofs. We show strong evidence that under neutrality our approaches can recover a constant (and correct) *α* along the genome as well as the past variation of the population size. We further predict that, while selective processes may preferentially occur in coding regions or regulatory potentially non-coding regions, local variations in *α* (as a consequence of sweepstake events) should be indifferent to the genomic functionality (coding or non-coding). Hence, we suggest that current sequence simulators [7, 36] could be extended to include the aforementioned factors and *de facto* facilitate the development of machine learning approaches.

Our study is unique in developing a state-of-the-art SMC approach and demonstrating that computational and mathematical problems can be overcome by deep learning (here GNN) approaches. The GNN*coal* approach is, in principle, not limited to the *β*-coalescent, and should work for other multiple merger models (*e.g.*, Dirac coalescents [28]) with the appropriate training. Furthermore, our SM*β*C approach is the first step to build a full genome method with an underlying model accounting for positive selection. In the future, further implementations may be added for a more realistic approach. The *α* parameter should be varying along the genome (as a hidden state), as the recombination rate in the iSMC [5]. This would allow to account for the local effect of strong and weak selection [1]. The effect of the *α* parameter could be also changing through time to better model the non uniform occurrence of multiple merger events through time. Although it is mathematically correct to have *α* as a constant in time, it is erroneous in practice (Figure S2). We speculate that those additional features will allow to accurately model and infer multiple merger events, variation of population size, and selection at each position on the genome. We believe that deep learning approaches could also be improved to recover more complex scenarios, providing in depth development on the structure of the graph neural networks, for example, by accounting for more features. At last, further investigation are required to make progress in the interpretability of the GNN methods, namely which statistics and convolution of statistics are used by GNN*coal* to infer which parameters.

As our approaches are the first of their kind, we chose to restrain our study to haploid models of *β* and Kingman coalescent as a proof of principle. However, the GNN*coal* and SM*β*C approaches can be extended to higher ploidy levels. Diploid versions of the haploid reproduction models whose genealogies are given by the *β*-coalescent lead to slightly different MMC coalescent models which can exhibit simultaneous multiple mergers [8, 10]. Thus, our GNN approach should be directly applicable when trained on these diploid models which are implemented in *msprime* [7]. However, to adjust the SM*β*C approach would be somewhat more cumbersome (but doable), since we would need to extend the underlying HMM to account for simultaneous multiple mergers. We emphasise that while there is growing evidence that MMC models produce better fitting genealogies for various species [33], there is ongoing discussions about which mathematical models are better suited to which species (for example see [3] for cod). We advocate that the lifecycle and various ecological factors determine whether a haploid or diploid MMC model can be chosen. On the one hand, a diploid MMC model is likely realistic if the species has a diploid life-cycle and balanced sex-ratio, so that multiple merger events do indeed happen in both sexes. On the other hand, if species are mostly haploid or clonal/asexual during their life-cycle (with periodically one short diploid phase for sexual reproduction) or exhibit strongly imbalanced sex-ratio, a haploid MMC model may be better suited. In their current form, our approaches are applicable to data from species with the latter characteristics such as many fungal and micro-parasites of plants and animals (including humans) as well as invertebrates (*e.g. Daphnia* or aphids) which undergo several clonal or parthenogenetic phases of reproduction (and one short sexual phase) per year. This represents a non-negligible set of study organisms which are of importance for medicine and agriculture [94].

Our results on inferred ARGs stress the need for improving ARG inference [15]. Thanks to the SMC we are close to model the ARG allowing to infer demographic history, selection and specific reproductive mechanism. Moreover, the comparison of deep learning approaches with model driven *ad hoc* SMC methods may have the potential to help us solve ongoing challenges in the field. These include simultaneously inferring and accounting for recombination, variation of population size, different type of selection, population structure and the variation of the mutation and recombination rate along the genome. These issues have puzzled theoreticians and statisticians since the dawn of population genetics [44].

On a final note, as environmental changes hit us all, we suggest that decreasing the computer and power resources needed to perform DL/ GNN analyses should be attempted [82]. Based on our study, we suggest that population genetics DL methods could be built as a two step process: 1) inferring ARGs, and 2) inferring demography and selection based on the ARGs. We speculate that general training sets based on ARGs could be build and be widely applicable for inference across many species with different life cycles and life history traits, while the inference of ARGs could be undertaken by complementary deep learning or Hidden Markov methods.

## Data availability

Code used to generate the simulated data for analysis, training and validation alongside (trained) deep learning models can be found at https://github.com/kevinkorfmann/ GNNcoal and https://github.com/kevinkorfmann/GNNcoal-analysis. Code for SMC approaches used in this manuscript are available in the R package eSMC2 https://github.com/TPPSellinger/eSMC2. msprime and its documentation can be found: https://tskit.dev/msprime/docs/stable/quickstart.html.

## Acknowledgments

This work was supported by the BMBF-funded de.NBI Cloud within the German Network for Bioinformatics Infrastructure (de.NBI) (031A532B, 031A533A, 031A533B, 031A534A, 031A535A, 031A537A, 031A537B, 031A537C, 031A537D, 031A538A). KK is supported by a grant from the Deutsche Forschungsgemeinschaft (DFG) through the TUM International Graduate School of Science and Engineering (IGSSE), GSC 81, within the project GENOMIE QADOP. TS is supported by the Austrian Science Fund (project no. TAI 151-B). AT acknowledges funding from the DFG grant TE809/1-4 (project 254587930) and TE809/7-1 (project 317616126). FF and AT acknowledge funding from the DFG Priority Program SPP1590 on “Probabilistic Structures in Evolution”. MF and AT acknowledge the support from the Imperial College - TUM Partnership award.

