## Supplementary Figures and Tables for "Simultaneous Inference of Past Demography and Selection from the Ancestral Recombination Graph under the Beta Coalescent"

### 1 Supplementary Figures

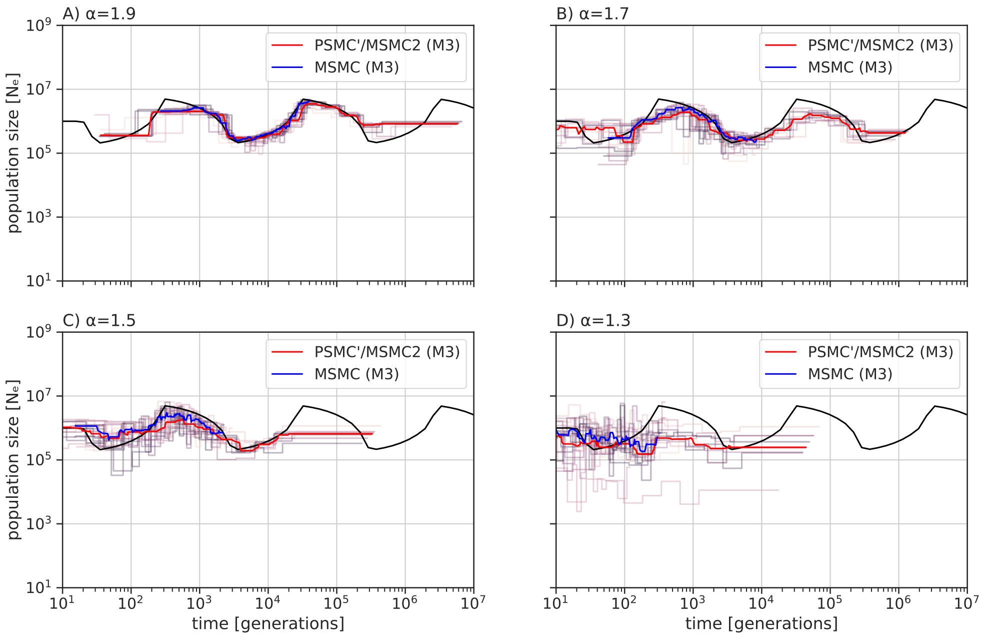


**Fig. S1.** Estimated demographic history by MSMC (blue) and MSMC2 (red) from the true ARG using 10 sequences of 100 Mb when population undergoes a sawtooth demographic scenario (black) for different $\alpha$ values: A) 1.9, B) 1.7, C) 1.5,and D) 1.3. The estimated population size is corrected in order to mask the scaling difference between the Kingman coalescent and the Beta coalescent ($i.e.$ $Ne = (\frac{\mu_{estimated}}{\mu_{real}})/scale)^{\frac{1}{(\alpha-1)}}$, where $m = 1 + \frac{1}{2^{\alpha-1}\times(\alpha-1)}$, $scale = \frac{(m^{\alpha})}{(\alpha*\beta(2-\alpha,\alpha))}$ and $\mu_{\text{estimated}} = \frac{\theta}{\left(2\cdot\sum_{i=1}^{n_{\text{ind}}-1}\frac{1}{i}\right)\cdot L}$). The recombination and mutation rate are set to $1 \times 10^{-8}$ per generation per bp.

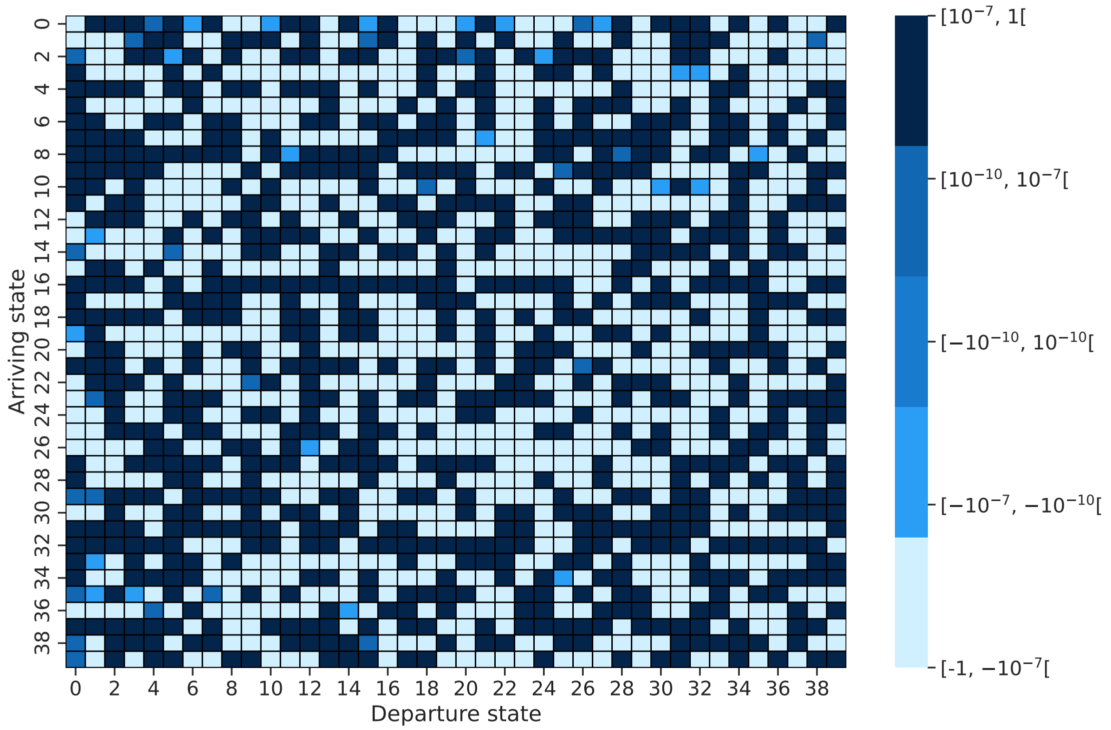


**Fig. S2.** Difference between the observed transition matrix and the theoretical prediction under the eSMC2 under a constant population size under the Kingman Coalescent

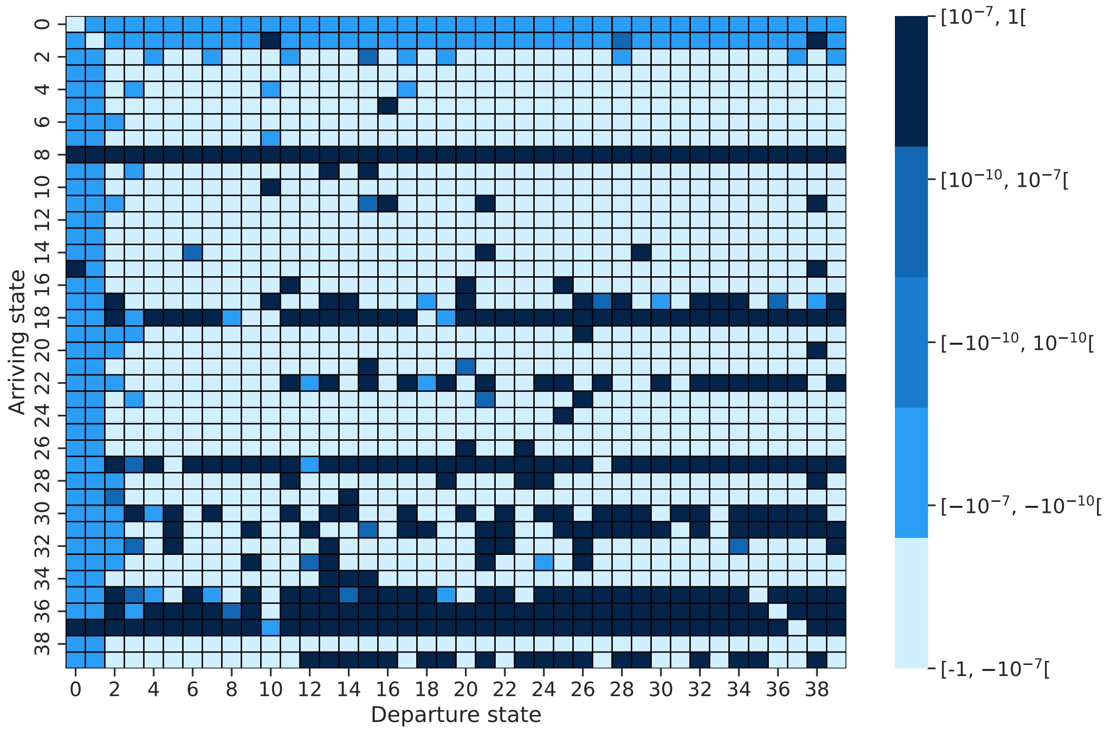


**Fig. S3.** Difference between the observed transition matrix and the theoretical prediction under the eSMC2 under a constant population size under the Beta Coalescent with $\alpha = 1.3$

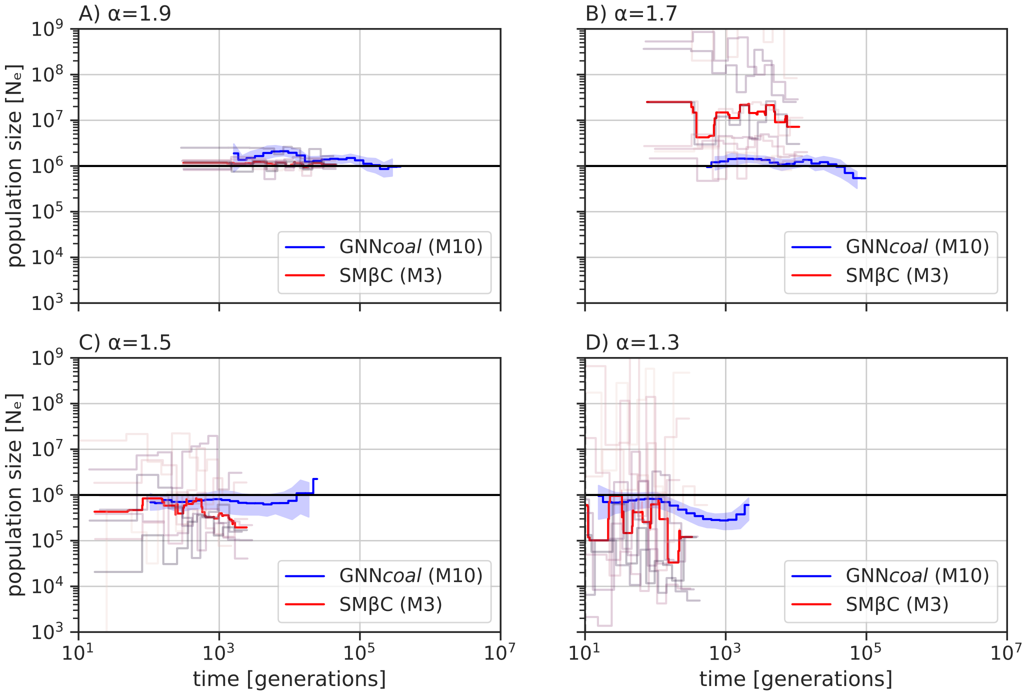


**Fig. S4. Best-case convergence estimations of SM$\beta$C and GNN*coal* under a Beta coalescent and constant population size.** Estimations of past demographic history by SM$\beta$C using 10 sequences and 100 Mb in red (median) and by GNN using 10 sequences and 500 trees in blue (mean and CI95) when population undergoes a "constant" demographic scenario (black) under 4 different $\alpha$ values 1.9,1.7,1.5 and 1.3, respectively in A),B),C) and D). The recombination and mutation rate are set to $1 \times 10^{-8}$ per generation per bp.

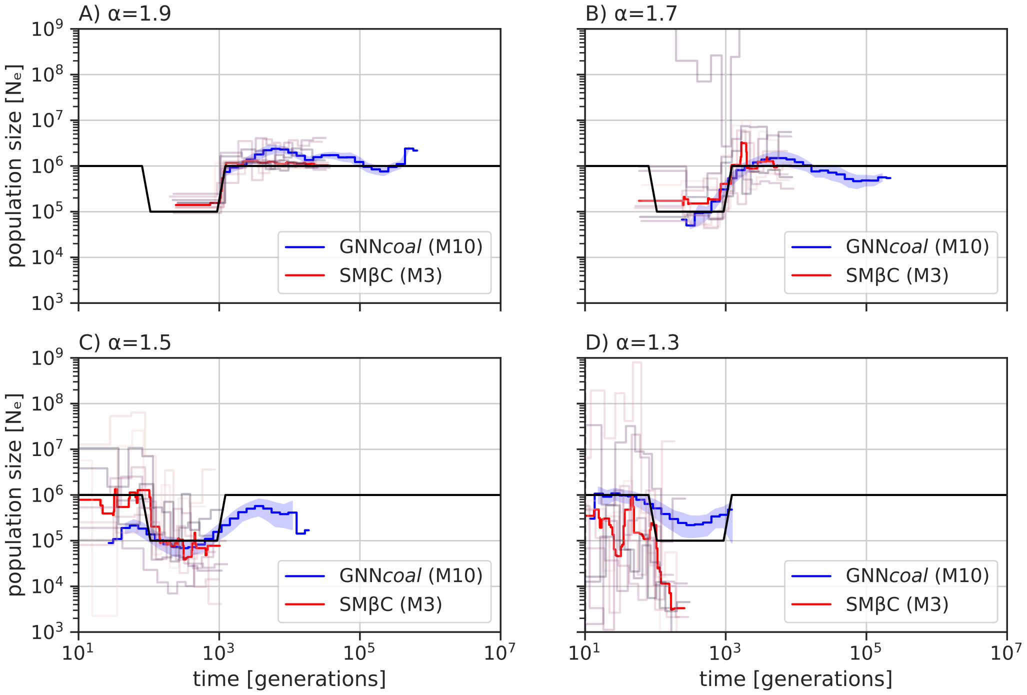


**Fig. S5. Best-case convergence estimations of SM$\beta$C and GNN*coal* under a Beta coalescent when the population undergoes a "bottleneck".** Estimations of past demographic history by SM$\beta$C using 10 sequences and 100 Mb in red (median) and by GNN using 10 sequences and 500 trees in blue (mean and CI95) when population undergoes a "bottleneck" demographic scenario (black) under 4 different $\alpha$ values A) 1.9, B) 1.7, c) 1.5 and D) 1.3. The recombination and mutation rate are set to $1 \times 10^{-8}$ per generation per bp.

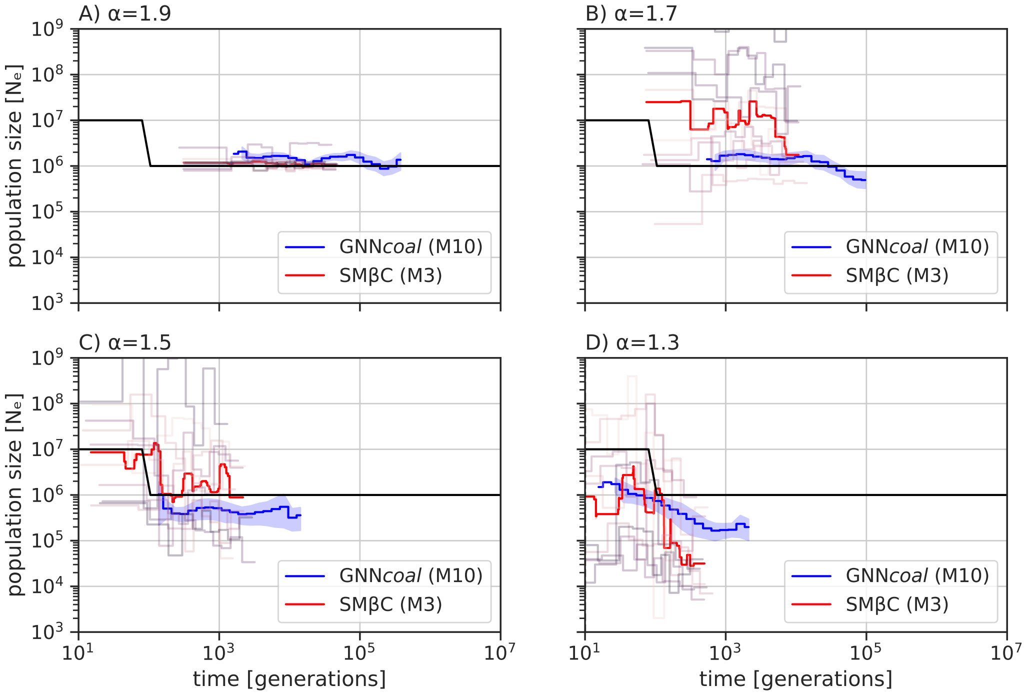


**Fig. S6. Best-case convergence estimations of SM$\beta$C and GNN*coal* under a Beta coalescent when population undergoes a sudden increase in size.** Estimations of past demographic history by SM$\beta$C using 10 sequences and 100 Mb in red (median) and by GNN using 10 sequences and 500 trees in blue (mean and CI95) when population undergoes a "increase" demographic scenario (black) under 4 different $\alpha$ values A) 1.9, B) 1.7, c) 1.5 and D) 1.3. The recombination and mutation rate are set to $1 \times 10^{-8}$ per generation per bp.

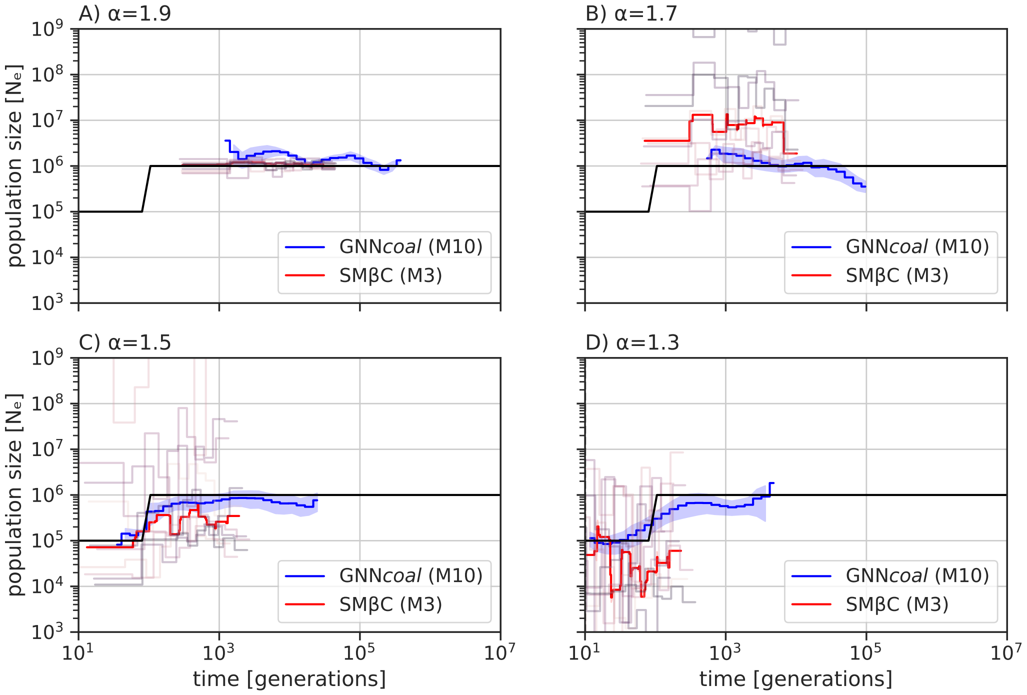


**Fig. S7. Best-case convergence estimations of SM$\beta$C and GNN*coal* under a Beta coalescent when population undergoes a sudden decrease in size.** Estimations of past demographic history by SM$\beta$C using 10 sequences of 100 Mb in red (median) and by GNN using 10 sequences and 500 trees in blue (mean and CI95) when population undergoes a "decrease" demographic scenario (black) under 4 different $\alpha$ values A) 1.9, B) 1.7, c) 1.5 and D) 1.3. The recombination and mutation rate are set to $1 \times 10^{-8}$ per generation per bp.

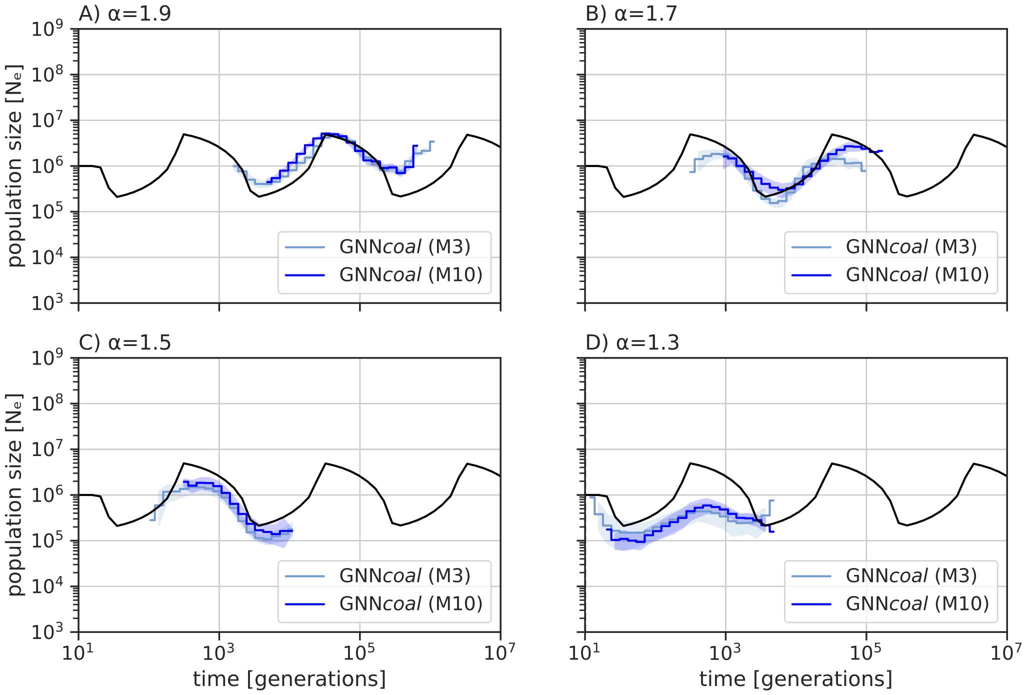


**Fig. S8. Best-case convergence estimations of two GNNs under a Beta coalescent when population undergoes a "Sawtooth" scenario.** Estimations of past demographic history by GNN*coal* using 10 sequences in blue and by a GNN*coal* using 3 sequences in ice blue when population undergoes a "Sawtooth" demographic scenario (black) under 4 different $\alpha$ values A) 1.9, B) 1.7, c) 1.5 and D) 1.3 (mean and CI95). The recombination and mutation rate are set to $1 \times 10^{-8}$ per generation per bp.

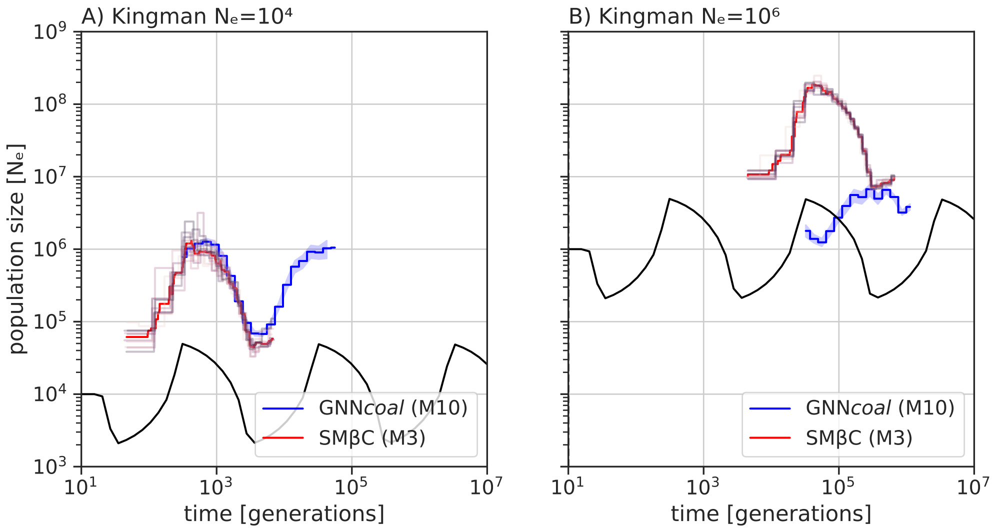


**Fig. S9. Best-case convergence estimations of SM$\beta$C and GNN*coal* under a Kingman coalescent when population undergoes a "Sawtooth" scenario.** Estimations of past demographic history by SM$\beta$C using 10 sequences and 10 Mb in red (median) and by GNN using 10 sequences and 500 trees in blue (mean and CI95) when population undergoes a "sawtooth" demographic scenario (black) under 2 different population sizes A) $N_e = 10^4$ and B) $N_e = 10^6$. The recombination and mutation rate are set to $1 \times 10^{-8}$ per generation per bp.

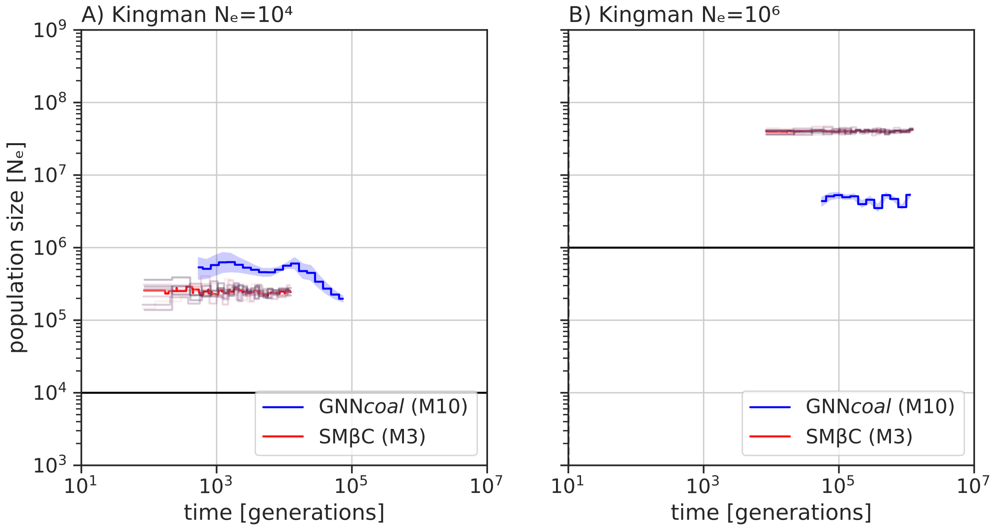


**Fig. S10. Best-case convergence estimations of SM$\beta$C and GNN*coal* under a Kingman coalescent when population stays constant.** Estimations of past demographic history by SM$\beta$C using 10 sequences and 10 Mb in red (median) and by GNN using 10 sequences and 500 trees in blue (mean and CI95) when population undergoes a "constant" demographic scenario (black) under 2 different population sizes A) $N_e = 10^4$ and B) $N_e = 10^6$. The recombination and mutation rate are set to $1 \times 10^{-8}$ per generation per bp.

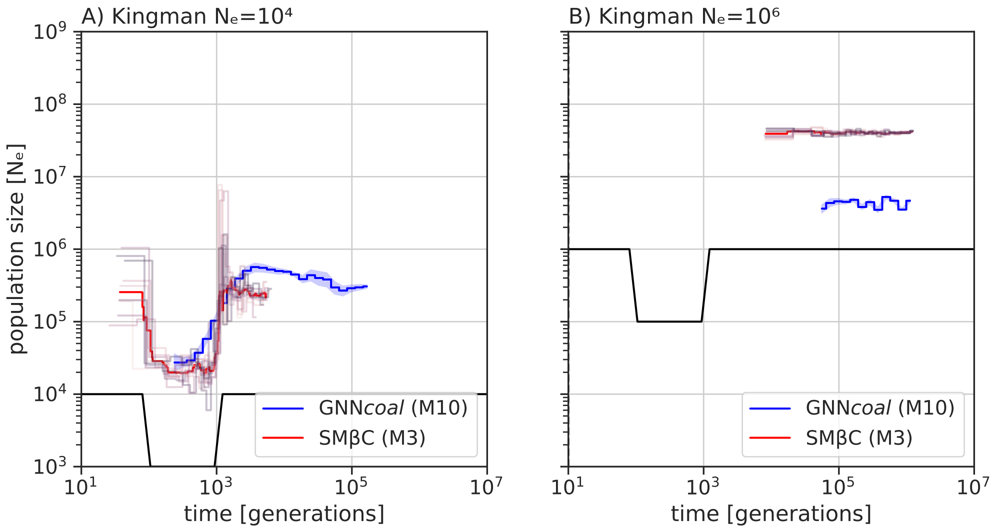


**Fig. S11. Best-case convergence estimations of SM$\beta$C and GNN*coal* under a Kingman coalescent when population undergoes a sudden "bottleneck".** Estimations of past demographic history by SM$\beta$C using 10 sequences and 10 Mb in red (median) and by GNN using 10 sequences and 500 trees in blue (mean and CI95) when population undergoes a "bottleneck" demographic scenario (black) under 2 different population sizes A) $N_e = 10^4$ and B) $N_e = 10^6$. The recombination and mutation rate are set to $1 \times 10^{-8}$ per generation per bp.

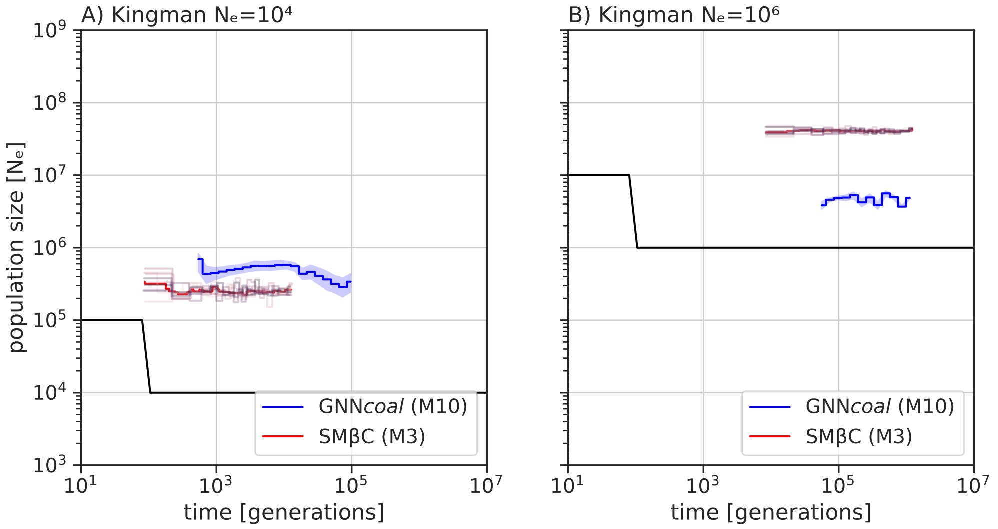


**Fig. S12. Best-case convergence estimations of SM$\beta$C and GNN*coal* under a Kingman coalescent when population undergoes a sudden increase in size.** Estimations of past demographic history by SM$\beta$C using 10 sequences and 10 Mb in red (median) and by GNN using 10 sequences and 500 trees in blue (mean and CI95) when population undergoes a "increase" demographic scenario (black) under 2 different population sizes A) $N_e = 10^4$ and B) $N_e = 10^6$. The recombination and mutation rate are set to $1 \times 10^{-8}$ per generation per bp.

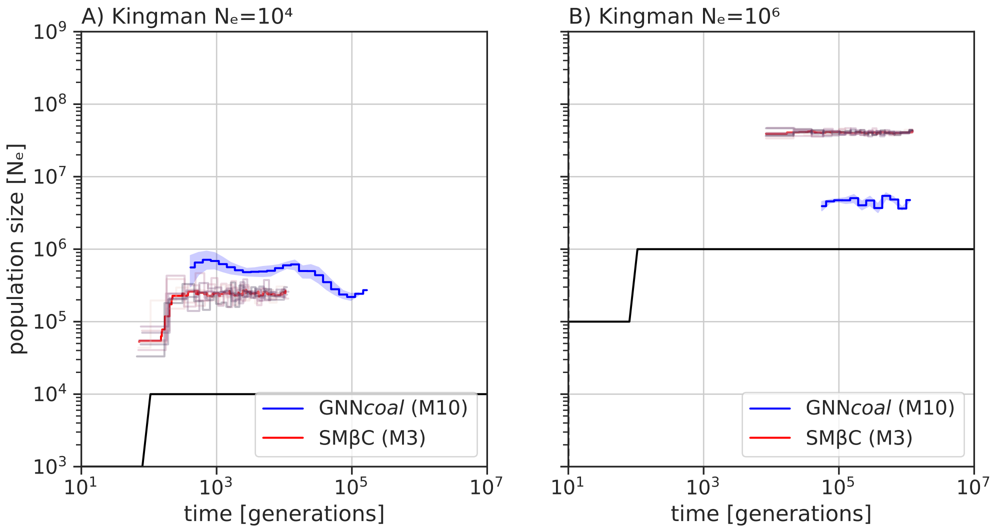


**Fig. S13. Best-case convergence estimations of SM$\beta$C and GNN*coal* under a Kingman coalescent when population undergoes a sudden decrease in size.** Estimations of past demographic history by SM$\beta$C using 10 sequences and 10 Mb in red (median) and by GNN using 10 sequences and 500 trees in blue (mean and CI95) when population undergoes a "decrease" demographic scenario (black) under 2 different population sizes A) $N_e = 10^4$ and B) $N_e = 10^6$. The recombination and mutation rate are set to $1 \times 10^{-8}$ per generation per bp.

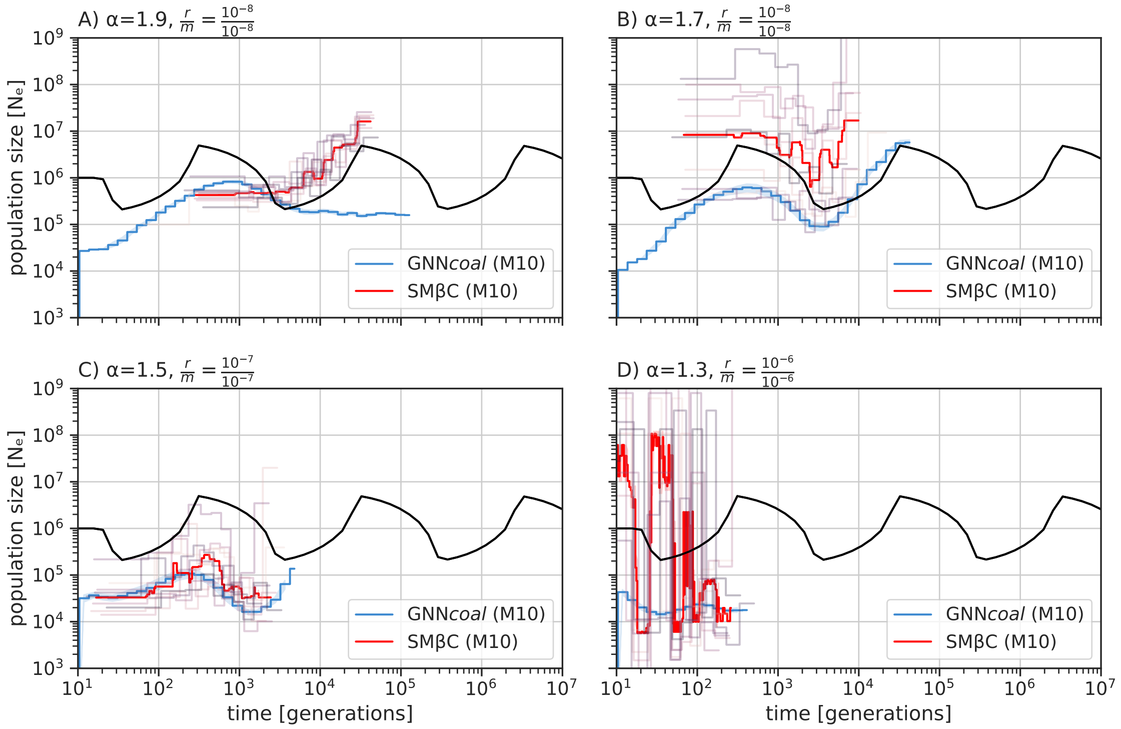


**Fig. S14. Demographic inference estimations of SM$\beta$C and GNN*coal* using ARG-weaver output under a sawtooth scenario.**. Estimations of the ARG is first performed by ARGweaver using 10 sequences of10 Mb. Estimations of past demographic history is then performed on the inferred ARG by SM$\beta$C in red (median) and by GNN using 10 sequences and 500 trees in blue (mean and CI95) when population undergoes a "sawtooth" demographic scenario (black). The recombination and mutation rate per generation per bp are set to $1 \times 10^{-8}$ for $\alpha$ equal to 1.9 and 1.7 , $1 \times 10^{-7}$ for $\alpha$ equal to 1.5 and set to $1 \times 10^{-6}$ for $\alpha$ equal to 1.3.

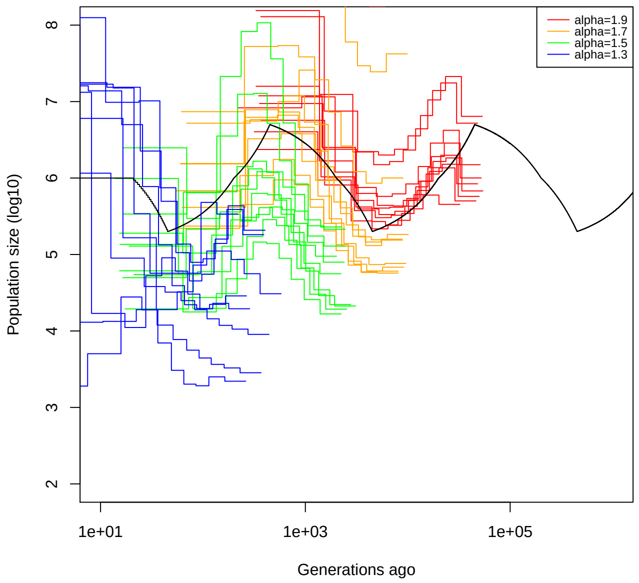


**Fig. S15. Demographic inference estimations of SM$\beta$C on simulated data under a sawtooth demographic scenario**.Estimated population size by SM$\beta$C from simulated sequence data under a sawtooth demographic scenario (black) and the Beta coalescent. Estimations of past demographic history by SM$\beta$C using 10 sequences of 10 Mb under different $\alpha$ values ( 1.9 in red, 1.7 in orange, 1.5 in green and 1.3 in blue. The recombination and mutation rate per generation per bp are set to $1 \times 10^{-8}$ for $\alpha$ equal to 1.9 and 1.7 , $1 \times 10^{-7}$ for $\alpha$ equal to 1.5 and set to $1 \times 10^{-6}$ for $\alpha$ equal to 1.3 .

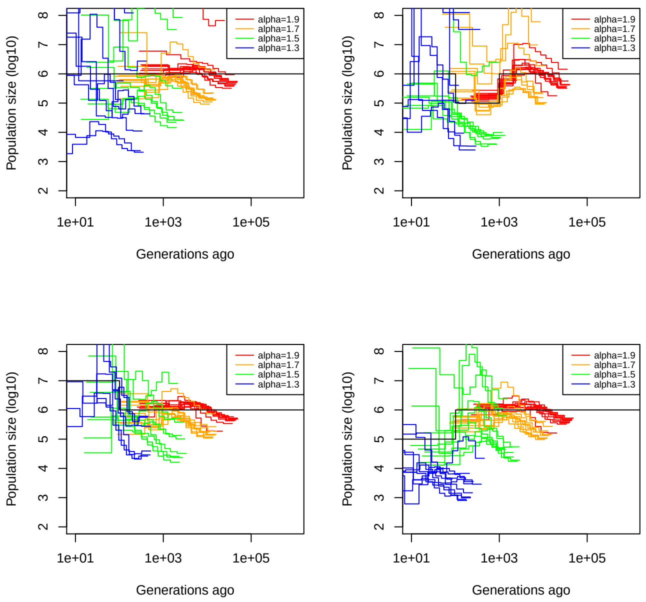


**Fig. S16. Demographic inference estimations of SM$\beta$C on simulated data under 4 difference demographic scenarios.**Estimated population size by SM$\beta$C from simulated sequence data under 4 demographic scenarios (black) (Constant population size in A, Bottleneck in B, Sudden increase of population size in C, and sudden decrease in D). Sequences are simulated under the Beta coalescent. Estimations of past demographic history by SM$\beta$C using 10 sequences of 10 Mb under different $\alpha$ values ( 1.9 in red, 1.7 in orange, 1.5 in green and 1.3 in blue. The recombination and mutation rate per generation per bp are set to $1 \times 10^{-8}$ for $\alpha$ equal to 1.9 and 1.7 , $1 \times 10^{-7}$ for $\alpha$ equal to 1.5 and set to $1 \times 10^{-6}$ for $\alpha$ equal to 1.3 .

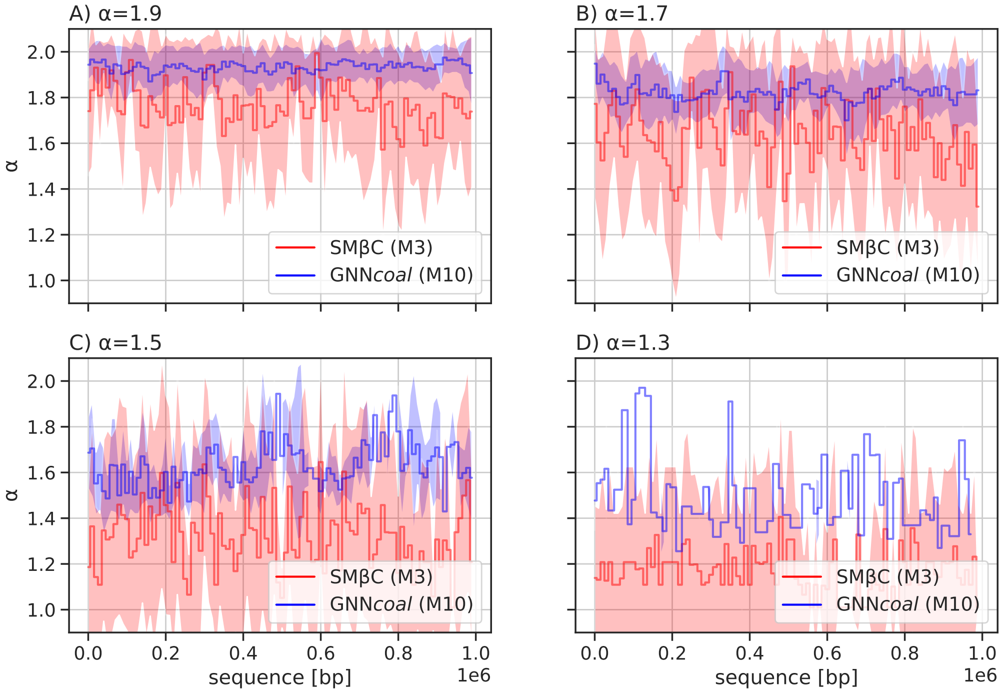


**Fig. S17. Averaged estimations of alpha by the GNN*coal* approach and the SM$\beta$C along the sequence** Estimations of $\alpha$ by SM$\beta$C using 20 sequences 1 Mb in red and blue by GNN*coal* using 10 sequences and in blue under 4 different $\alpha$ values 1.9,1.7,1.5 and 1.3 for a constant demography of size $10^6$ in A),B),C) and D) (mean and standard deviation for both methods). The GNN*coal* used at most 20 coalescent trees per batch along the sequence. Both methods used a window size of $10^3$ bp. The recombination and mutation rate are set to $1 \times 10^{-8}$ per generation per bp.

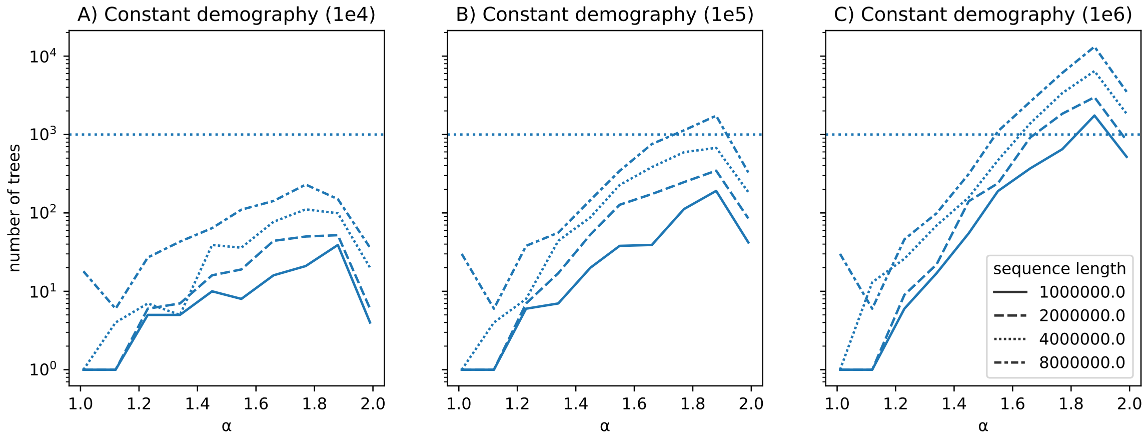


**Fig. S18. Tree sparsity with increasing $\alpha$.** Simulations of the $\beta$-coalescent to capture number of genealogies as a function of the multiple merger strength ($\alpha$ parameter of the $\beta$-coalescent). A constant population size was set ranging from $10^4$ (A) to $10^6$ (C) with one simulations per $\alpha$ at positions for $\alpha$ equal to 1.01, 1.11, 1.22, 1.33, 1.44, 1.55, 1.66, 1.77, 1.88 and 1.99. The threshold of 1000 genealogies is marked as horizontal line.

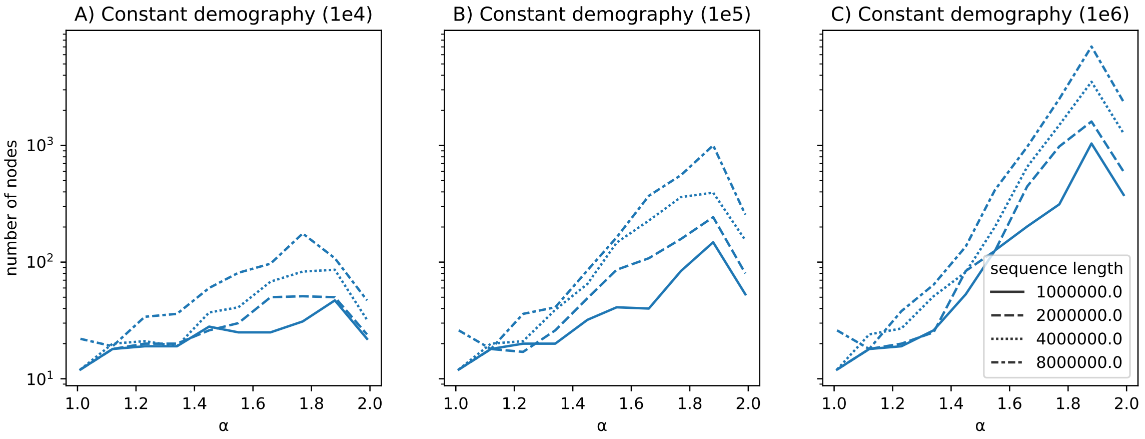


**Fig. S19. Node (ancestor+leaf) sparsity with increasing $\alpha$.** Simulations of the $\beta$-coalescent to capture number of nodes as a function of the multiple merger strength ($\alpha$ parameter of the $\beta$-coalescent). A constant population size was set ranging from $10^4$ (A) to $10^6$ (C) with one simulation per $\alpha$ at positions for $\alpha$ equal to 1.01, 1.11, 1.22, 1.33, 1.44, 1.55, 1.66, 1.77, 1.88 and 1.99.

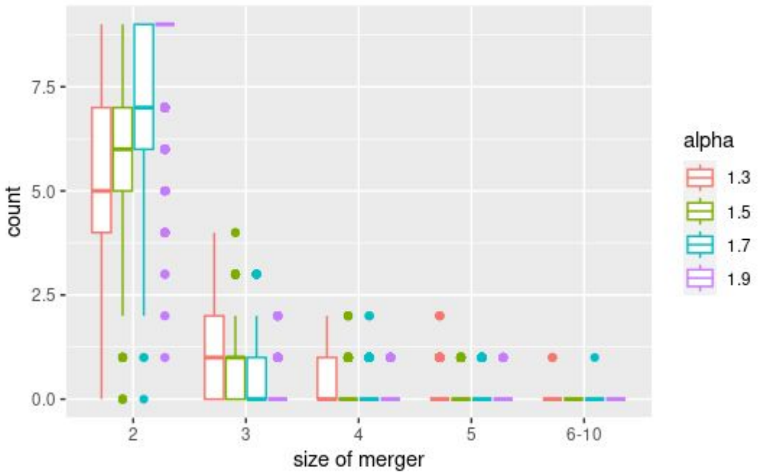


**Fig. S20. Counts of multiple merger sizes with regard to $\alpha$.** Simulations of the $\beta$-coalescent to capture counts and types of multiple mergers based on 10,000 repetitions for sample size of 10 haploid individuals (modelling a single locus, no recombination).

### 2 Supplementary Tables

| scenario | True $\alpha$ | $\alpha^*$:SM$\beta$C,M=3 | $\alpha^*$:SM$\beta$C,M=4 | $\alpha^*$ : GNN, M=3 | $\alpha^*$ : GNN, M=10 |
|---|---|---|---|---|---|
| Constant | 1.9 | 1.88 (0.04) | 1.84 (0.1) | 1.90 (0.03) | 1.88 (0.03) |
| Bottleneck | 1.9 | 1.86 (0.07) | 1.82 (0.16) | 1.91 (0.03) | 1.91 (0.02) |
| Increase | 1.9 | 1.88 (0.05) | 1.86 (0.12) | 1.91 (0.03) | 1.88 (0.02) |
| Decrease | 1.9 | 1.89 (0.02) | 1.88 (0.07) | 1.93 (0.03) | 1.88 (0.04) |
| Sawtooth | 1.9 | 1.84 (0.06) | 1.74 (0.11) | 1.96 (0.02) | 1.94 (0.02) |
| Constant | 1.7 | 1.53 (0.12) | 1.63 (0.09) | 1.78 (0.03) | 1.77 (0.03) |
| Bottleneck | 1.7 | 1.67 (0.14) | 1.57 (0.09) | 1.78 (0.04) | 1.76 (0.03) |
| Increase | 1.7 | 1.57 (0.14) | 1.63 (0.17) | 1.77 (0.01) | 1.74 (0.03) |
| Decrease | 1.7 | 1.54 (0.15) | 1.57 (0.14) | 1.80 (0.04) | 1.78 (0.04) |
| Sawtooth | 1.7 | 1.67 (0.14) | 1.60 (0.19) | 1.81 (0.02) | 1.74 (0.03) |
| Constant | 1.5 | 1.58 (0.10) | 1.56 (0.14) | 1.61 (0.02) | 1.56 (0.02) |
| Bottleneck | 1.5 | 1.53 (0.11) | 1.55 (0.19) | 1.58 (0.02) | 1.53 (0.03) |
| Increase | 1.5 | 1.50 (0.12) | 1.52 (0.14) | 1.64 (0.05) | 1.61 (0.03) |
| Decrease | 1.5 | 1.57 (0.17) | 1.55 (0.14) | 1.60 (0.03) | 1.55 (0.03) |
| Sawtooth | 1.5 | 1.47 (0.11) | 1.55 (0.19) | 1.61 (0.04) | 1.56 (0.02) |
| Constant | 1.3 | 1.40 (0.13) | 1.45 (0.14) | 1.39 (0.03) | 1.33 (0.03) |
| Bottleneck | 1.3 | 1.43 (0.13) | 1.48 (0.20) | 1.34 (0.02) | 1.28 (0.04) |
| Increase | 1.3 | 1.45 (0.14) | 1.51 (0.24) | 1.42 (0.02) | 1.38 (0.02) |
| Decrease | 1.3 | 1.44 (0.16) | 1.50 (0.12) | 1.4 (0.02) | 1.30 (0.03) |
| Sawtooth | 1.3 | 1.39 (0.09) | 1.39 (0.07) | 1.41 (0.03) | 1.36 (0.05) |

Table S1: Average estimated values of $\alpha$ by SM$\beta$C and the GNN*coal* approach over ten repetitions using the true ARG of 10 sequences of 100 Mb with recombination and mutation rate set to $1 \times 10^{-8}$ per generation per bp under a Beta coalescent process (with different $\alpha$ parameter). The analysis were run on five different demographic scenarios (Constant population size, Bottleneck, Sudden increase, Sudden decrease and a Sawtooth demography). The standard deviation is indicated in brackets. These results complement the demographic estimates from Figures 4 and 5, as well as Supplementary Figure 4 to 7

| scenario | True $\alpha$ | $\alpha^*$:SM$\beta$C,M=3 | $\alpha^*$:SM$\beta$C,M=4 |
|---|---|---|---|
| Constant | 1.7 | 1.64 (0.12) | 1.56 (0.11) |
| Bottleneck | 1.7 | 1.59 (0.15) | 1.55 (0.12) |
| Increase | 1.7 | 1.65 (0.10) | 1.61 (0.15) |
| Decrease | 1.7 | 1.71 (0.10) | 1.67 (0.12) |
| Sawtooth | 1.7 | 1.60 (0.18) | 1.51 (0.20) |
| Constant | 1.5 | 1.46 (0.11) | 1.51 (0.16) |
| Bottleneck | 1.5 | 1.50 (0.14) | 1.45 (0.16) |
| Increase | 1.5 | 1.56 (0.13) | 1.47 (0.15) |
| Decrease | 1.5 | 1.51 (0.11) | 1.47 (0.12) |
| Sawtooth | 1.5 | 1.40 (0.13) | 1.44 (0.17) |
| Constant | 1.3 | 1.38 (0.15) | 1.37 (0.08) |
| Bottleneck | 1.3 | 1.34 (0.10) | 1.34 (0.06) |
| Increase | 1.3 | 1.34 (0.08) | 1.46 (0.17) |
| Decrease | 1.3 | 1.41 (0.15) | 1.44 (0.09) |
| Sawtooth | 1.3 | 1.35 (0.07) | 1.34 (0.05) |

Table S2: Average estimated values of $\alpha$ by SM$\beta$C over ten repetitions using the true ARG of 10 sequences of 100 Mb with recombination and mutation rate set to $5 \times 10^{-8}$ for $\alpha = 1.7$ per generation per bp and $5 \times 10^{-7}$ for $\alpha$ equal to 1.5 and 1.3 under a Beta coalescent process (with different $\alpha$ values). The coefficient of variation is indicated in brackets.

| scenario | True $\alpha$ | $\alpha^*$:SM$\beta$C,M=3 | $\alpha^*$:SM$\beta$C,M=4 | $\alpha^*$ : GNN, M=3 | $\alpha^*$ : GNN, M=10 |
|---|---|---|---|---|---|
| Constant | 2 | 1.98 (0.023) | 1.97 (0.023) | 1.99 (0.002) | 1.99 (0.005) |
| Sawtooth | 2 | 1.96 (0.027) | 1.87 (0.064) | 1.98 (0.003) | 1.99 (0.005) |
| Bottleneck | 2 | 1.96 (0.047) | 1.96 (0.04) | 1.99 (0.002) | 1.99 (0.003) |
| Decrease | 2 | 1.98 (0.022) | 1.99 (0.012) | 1.98 (0.009) | 1.99 (0.005) |
| Increase | 2 | 1.97 (0.020) | 1.97 (0.025) | 1.99 (0.002) | 1.99 (0.004) |

Table S3: Average estimated values of $\alpha$ by SM$\beta$C and the GNN*coal* approach over ten repetitions using the true ARG of 10 sequences of 100 Mb for SM$\beta$C and at 500 trees for the GNN*coal* with recombination and mutation rate set to $1 \times 10^{-8}$ per generation per bp under a Beta coalescent process (with different $\alpha$ parameter) with $N_e = 10^4$ at generation 0. The standard deviation is indicated in brackets. These results complement the estimations of demography from Supplementary Figure 9-13 .

| scenario | True $\alpha$ | $\alpha^*$:SM$\beta$C,M=3 | $\alpha^*$:SM$\beta$C,M=4 | $\alpha^*$ : GNN, M=3 | $\alpha^*$ : GNN, M=10 |
|---|---|---|---|---|---|
| Sawtooth | 1.9 | 1.77 (0.05) | 1.86 (0.05) | 1.44 (0.005) | 1.98 (0.001) |
| Sawtooth | 1.7 | 1.56 (0.13) | 1.79 (0.1) | 1.44 (0.002) | 1.98 (0.002) |
| Sawtooth | 1.5 | 1.66 (0.11) | 1.92 (0.07) | 1.40 (0.003) | 1.97 (0.002) |
| Sawtooth | 1.3 | 1.41 (0.13) | 1.53 (0.20) | 1.33 (0.004) | 1.78 (0.027) |

Table S4: Average estimated values of $\alpha$ by SM$\beta$C and the GNN*coal* approach using ARG inferred by ARGweaver over ten repetitions using 10 simulated sequences of 10 Mb with recombination and mutation rate set to $1 \times 10^{-8}$ per generation per bp under a Beta coalescent process (with different $\alpha$ parameter). The analysis were run under Sawtooth demographic scenario. The standard deviation is indicated in brackets. These results complement the demographic estimates from Supplementary Figures 14
