## Supplementary Text S1 for "Simultaneous Inference of Past Demography and Selection from the Ancestral Recombination Graph under the Beta Coalescent"

### **Supplementary Text S1:** The Sequentially Markovian $\beta$ Coalescent

Kevin Korfmann, Thibaut Sellinger, Fabian Freund,
Matteo Fumagalli, Aurélien Tellier

#### 1 SM$\beta$C

The Sequentially Markovian $\beta$ Coalescent is a Hidden Markov Model based on the Multiple Sequentially Markovian Coalescent (MSMC) where Multiple Merger event are allowed to occur under the $\beta$ coalescent.

To define our Hidden Markov Model (HMM) we need to define :

- Hidden States
- The signal (observed data)
- A Transition matrix (Probability of passing from one state to another)
- An Emission matrix (Probability of observing the signal conditional to the hidden state)
- An Initial probability (Probability of hidden states at the first position of the sequence)

##### 1.1 Notations and Assumptions

We here define the different notations used and their meaning:

- $r$ : recombination rate per nucleotide
- $\mu$ : Mutation rate per nucleotide
- $u$ : recombination time, follows a continuous uniform distribution between 0 and first coalescent time.
- $\xi_t$ : Scaled population size at time t ($N_t = \xi_t N_0$)
- $\chi_t = \xi_t^{\beta-1}$
- $M$ : Number of analyzed haploid sequences (or haploid individuals)
- $\beta$ : The multiple merger parameter

The model's assumptions are :

- $(\chi_t)_{t\geq 0}$ is piece-wise constant (intervals are specified in the following)

We first define the transition rates of the Beta $n$-coalescent [3]. The rate of transition from a state with $b$ lineages to $b-n+1$ lineages, i.e. a merger of $n$ lineages is

$$\lambda_{b,\beta,b-n+1} = \frac{B(n-\beta, b-n+\beta)}{\Gamma(2-\beta)\Gamma(\beta)}. \tag{1}$$

$$\Lambda_{b,\beta,b-n+1} = \frac{\binom{b}{n} B(n-\beta, b-n+\beta)}{\Gamma(2-\beta)\Gamma(\beta)}. \tag{2}$$

Thus, the total rate is

$$\lambda_{b,\beta} = \sum_{k=2}^{b} \frac{\binom{b}{k} B(k-\beta, b-k+\beta)}{\Gamma(2-\beta)\Gamma(\beta)} \tag{3}$$

Waiting times are exponentially distributed in the coalescent for population size constant in time. For time-varying population sizes, we define the time-changed $\Lambda$-$n$-coalescent as the (rescaled) genealogy limit from a Wright-Fisher type Cannings model with skewed offspring distributions as introduce in [5], which leads to a time-change waiting time for coalescence events: If a waiting time has rate $\lambda$ in the standard Beta $n$-coalescent (started at some time $t_0$), it has a waiting time density of

$$f(t) = \frac{\lambda}{\chi(t)} e^{-\int_{t_0}^{t} \frac{\lambda}{\chi(s)} ds}, \tag{4}$$

which follows as described in [2].

#### 1.2 Hidden States

Our hidden states at one position are the first coalescent time $t > 0$ at that position and which individuals $i := (i_1, \ldots, i_n)$ coalesce in the corresponding coalescence. A transition from coalescent time s to time t or a change in the index $i$ can only occur when a recombination happens.

#### 1.3 Observations

##### 1.3.1 The input is sequence Data

The observation signal is the comparison of the M analyzed sequences. Thus the signal is a series of number indicating the allelic state of the sequences at each position. For M=3, under the infinite site model hypothesis, only 4 different state can be observed along the sequence. All sequences are the same at this position (indicated by a 0), or one of the three sequences is different from the two other (indicated by i, if the sequence i is different from the two other).

##### 1.3.2 The input is an ancestral recombination graph

If the inputted signal is an ancestral recombination graph (ARG), it will first be processed to obtain a tree sequence (*i.e.* sequence of coalescent trees). Using the definition of hidden state (see above) the tree sequence is transformed into a sequence of hidden state.

#### 1.4 Transition Matrix

Five transitions are possible, we transition from $(s, j)$ to $(t, i)$. Here we assume that t and s are in interval time $\alpha$ and $\gamma$. At indices $i$ and $j$, $n$ and $m$ individuals coalesce. In addition, recombination occurs with probability :

$$P(rec|s) = 1 - e^{-rMs} \tag{5}$$

We assume that only a single recombination event can happen before a merger. A recombination event of one of $M$ lineages splits one ancestral lineage in two (backwards in time). The additional lineage is not yet described by the coalescent without the recombination event, we call this free. It can merge with any of the remaining $M$ lineages, but also with the second parental ancestral lineage (i.e. the second split lineage from the recombination event). The transition probabilities/rates conditional on the (known) behaviour of the other lineages are as described in [1, Sect. 5]: Conditional on the mergers of the $M$ other lineages, a binary merger of the "freed" lineage appears with rate $M\lambda_{M+1,\beta,M}$ and it joins an existing merger of $m$ lineages at some time $t$ with probability $1 - \frac{\lambda_{M+1,\beta,M+1-m+1}}{\lambda_{M,\beta,M-m+1}} = \frac{\lambda_{M+1,\beta,M+1-m}}{\lambda_{M,\beta,M-m+1}}$, where the second equation is due to the consistency of rates in $\Lambda$-$n$-coalescents. In the following, we derive conditional probabilities and/or conditional densities for certain events.

### 1.4.1 t<s

For this to happen, a recombination must occur before time $t$. The new number of individuals first coalescing is now $n = 2$, and the recombination event needs to affect one of these two individuals $i = \{i_1, i_2\}$ (which happens with probability $2/M$), splitting one lineage in two. Then, we just multiply the density of a binary merger of the free lineage with any of the other $M$ lineages in the time-changed coalescent, which is Eq. (4) with rate $M\lambda_{M+1,\beta,M}$ and the probability that the first merger is indeed merging $i$, which is $\frac{1}{M}$ (we pick the second lineage of $i$ at random from $M$ lineages):

$$f(t, i|s, j, u) = \frac{2\lambda_{M+1,\beta,M}}{M\chi_t} e^{-\int_u^t \frac{M\lambda_{M+1,\beta,M}}{\chi_v} dv} \tag{6}$$

### 1.4.2 t=s

Case 1: a non coalescing individual joins the coalescent event $j = \{j_1, \dots, j_n\}$. For this to happen, the recombination event must occur before time s in a non coalescing branch, which happens with probability $\frac{M-n}{M}$. Then, the newly split second ancestral lineage of $i$ need to not coalesce in a binary collision until time $s$, which equals $\exp(-\int_u^s \frac{M\lambda_{M+1,\beta,M}}{\chi_v} dv)$ (by integrating Eq. (4) with rate $M\lambda_{M+1,\beta,M}$). Finally, it then needs to join in the coalescent event $j$, which happens with probability $\frac{\lambda_{M+1,\beta,M+1-m}}{\lambda_{M,\beta,M-m+1}}$. This shows that

$$P(s, i|s, j, u) = \frac{(M-n)\lambda_{M+1,\beta,M+1-m}}{M\lambda_{M,\beta,M-m+1}} e^{-\int_u^s \frac{M\lambda_{M+1,\beta,M}}{\chi_v} dv} \tag{7}$$

for $i = j \cup \{i\}$.

Case 2: Recombination occurs in a coalescing individual $i \in j = \{j_1, \dots, j_n\}$ of a multiple merger event with $n > 2$ (happens with probability $\frac{1}{n}$). The new lineage then coalesces higher in time, i.e. it does neither coalesce in a binary merger before $s$ (as in case 1) nor in the collision at $s$ (which is the complementary event from case 1). As above, this leads to

$$P(s,i|s,j,u) = \frac{\lambda_{M+1,\beta,M+1-m+1}}{n\lambda_{M,\beta,M-m+1}} e^{-\int_u^s \frac{M\lambda_{M+1,\beta,M}}{\chi_v} dv} \tag{8}$$

for $i = j \setminus \{i\}$

Case 3: Nothing changes. This happens if a) there is no recombination event (so $u > s$), b) the lineage split makes a binary merger between the two lineages resulting from the split ("self-coalesce") before $s$, c) recombination splits a lineage merged at the coalescence event at $s$, but that the second ancestral lineage from the split joins the merger.
a) happens with probability 1 if $u > s$, b) has conditional density as in Eq. 6 without the factor $2/M$, integrating over $[u,s]$ yields the probability $(1-(1/M)\exp(-\int_u^t \frac{M\lambda_{M+1,\beta,M}}{\chi_v} dv)$ and c) follows as case 2, only we need the recombination event on a lineage already participating in the merger at time $s$ (so just replacing $M-n$ with $n$ in Eq. (8)).

### 1.4.3 t>s

For this to happen, a recombination must occur before time $s$ and break a coalescent event of only two individuals $(j_1, j_2)$ (w. probability $2/M$). Assume without restriction $j_1$ was affected by recombination. For (the new ancestral lineage of) $j_1$ to coalesce at time $t$, it must not coalesce until time s, then not coalesce in the former coalescent event and then the next coalescence event happens at time $t$. The next coalescence event can take any form and does not need to merge $j_1$. Additionally, we just need to keep track of the $M-1$ non-recombining lineages and $j_1$, since we are not conditioning on the behaviour after $s$ and thus both self-coalescence and the coalescence of the second split lineage (not $j_1$) can be ignored. Thus, to compute the conditional rate for merging into $i$, we first compute the probability that the new ancestral lineage representing $j_1$ in the new DNA segment does not coalesce until time $s$, given by Eq. (8) with $n = 2$, and multiply this by the conditional density for merging into any $i$ of M lineages afterwards. Thus, this is just Eq. (4) with rate $\lambda_{M,\beta}$ multiplied with $\frac{\lambda_{M,\beta,M-n+1}}{\lambda_{M,\beta}}$. This leads to

$$f(t,i|s,j,u) = \frac{2\lambda_{M+1,\beta,M}}{M\lambda_{M,\beta,M-1}} e^{-\int_u^s \frac{M\lambda_{M+1,\beta,M}}{\chi_v} dv} \frac{\lambda_{M,\beta,M-n+1}}{\chi_t} e^{-\int_s^t \frac{\lambda_{M,\beta}}{\chi_v} dv} \tag{9}$$

for $i = \{i_1, \ldots, i_n\}$.

##### 1.4.4 Full transition probability

$$p(t,i|s,j,u) = \begin{cases} (1-e^{-Mrs})\frac{2\lambda_{M+1,\beta,M}}{\chi_t M} e^{-\int_u^t \frac{M\lambda_{M+1,\beta,M}}{\chi_v} dv} & u < t < s \\ e^{-Mrs} + (1-e^{-Mrs})\left(\int_u^s \frac{e^{\int_u^k -\frac{M\lambda_{M+1,\beta,M}}{\chi_v} dv}}{\chi_k} dk + \frac{(M-n)\lambda_{(n+1),\beta,2}}{M\lambda_{(n+1),\beta,2}+\lambda_{(n+1),\beta,1}} e^{\int_u^t -\frac{M\lambda_{M+1,\beta,M}}{\chi_v} dv}\right) & t = s, m = n \\ (1-e^{-Mrs})\frac{(M-n)\lambda_{(n+1),\beta,1} e^{-\int_u^s \frac{M\lambda_{M+1,\beta,M}}{\chi_v} dv}}{M(\lambda_{(n+1),\beta,2}+\lambda_{(n+1),\beta,1})} & t = s, m = n+1 \\ (1-e^{-Mrs})\frac{1}{n}\frac{\lambda_{(n+1),\beta,2} e^{-\int_u^s \frac{M\lambda_{M+1,\beta,M}}{\chi_v} dv}}{s(\lambda_{(n+1),\beta,2}+\lambda_{(n+1),\beta,1})} & t = s, m+1 = n \\ (1-e^{-Mrs})\frac{\lambda_{M,\beta,(M-m+1)}}{\binom{M}{m}\chi_\alpha} e^{-\int_s^t \frac{\lambda_{M,\beta}}{\chi_v} dv} e^{-\int_u^s \frac{M\lambda_{M+1,\beta,M}}{\chi_v} dv} \frac{2\lambda_{(n+1),\beta,2}}{M(\lambda_{(n+1),\beta,2}+\lambda_{(n+1),\beta,1})} & t > s, i = l, j = k \end{cases} \tag{10}$$

As explained before, the state space must be finite. We therefore discretized time in k intervals. At one point the hidden state is $\alpha$ if $t \in [T_\alpha, T_{\alpha+1}]$, where $\alpha \in [0, (n-1)]$. We define $T_\alpha$ :

$$T_\alpha = \frac{-\ln(1-\frac{\alpha}{n})}{\lambda_{M,\beta}} \tag{11}$$

We therefore have:

$$p(\alpha, i|s, j) = \int_{T_\alpha}^{T_{\alpha+1}} p(t, i|s, j)dt \tag{12}$$

Note: Because time is discretized, if the first coalescent time is bigger than $T_{(n-1)}$, then all individual coalesce.

##### 1.4.5 Initial Probability

We use the equilibrium probability as initial probability while assuming m individual coalesce. The equilibrium probability is given by :

$$\begin{aligned} q_o(\alpha, i) &= \int_{T_\alpha}^{T_{\alpha+1}} \frac{\lambda_{M,\beta,(M-m+1)}}{\chi\alpha\binom{M}{m}} e^{-\int_0^t \frac{\lambda_{M,\beta}}{\chi v}dv} dt \\ &= \frac{\lambda_{M,\beta,(M-m+1)} e^{\sum_{\eta=0}^{\alpha-1} \frac{\lambda_{M,\beta}}{\chi\eta}\Delta_\eta}}{\binom{M}{m}\lambda_{M,\beta}} (1 - e^{-\Delta_\alpha \frac{\lambda_{M,\beta}}{\chi t}}) \end{aligned} \tag{13}$$

Where :

$$\Delta_\gamma = T_{\gamma+1} - T_\gamma \tag{14}$$

##### 1.4.6 Calculation of $t_{\gamma,j}$

Assuming $n$ individual coalesces.

$$\begin{aligned} t_{\gamma,j} = E[\text{Coalescent time}|\gamma, j] = \frac{E[\text{Coalescent time} \cap \gamma, j]}{P(\gamma, j)} &= \frac{\int_{T_\gamma}^{T_{\gamma+1}} t\lambda_{M,\beta,(M-n+1)} e^{-\int_0^t \frac{\lambda_{M,\beta}}{\chi v}dv}}{\binom{M}{n} q_0(\gamma, j)} dt \\ &= \frac{T_\gamma - T_{\gamma+1} e^{-\Delta_\gamma \frac{\lambda_{M,\beta}}{\chi\gamma}}}{(1 - e^{-\Delta_\gamma \frac{\lambda_{M,\beta}}{\chi\gamma}})} + \frac{\chi\gamma}{\lambda_{M,\beta}} \end{aligned} \tag{15}$$

Where :

$$\Delta_\gamma = T_{\gamma+1} - T_\gamma \tag{16}$$

We note that $t_{\gamma,j}$ is independent of $j$, thus $t_{\gamma,j} = t_\gamma$.

##### 1.4.7 Calculation of $p(\alpha, i|\gamma, j)$

$\alpha < \gamma$

We here calculate the transition probability from the state $\gamma$ to a time t in the time interval $\alpha$.

$$
\begin{aligned}
P(t,i|t_\gamma,j) &= \frac{P_\gamma}{t_\gamma}\int_0^t \frac{2\lambda_{M+1,\beta,M}}{\chi_v M} e^{-\int_u^t \frac{M\lambda_{M+1,\beta,M}}{\chi_v}dv} du \\
&= \frac{P_\gamma}{t_\gamma}(\sum_{\eta=0}^{\alpha-1}\int_{T_\eta}^{T_{\eta+1}} \frac{2\lambda_{M+1,\beta,M}}{\chi_v M} e^{-\int_u^t \frac{M\lambda_{M+1,\beta,M}}{\chi_v}dv} du + \int_{T_\alpha}^t \frac{2\lambda_{M+1,\beta,M}}{\chi_v M} e^{-\int_u^t \frac{M\lambda_{M+1,\beta,M}}{\chi_v}dv} du) \\
&= \frac{P_\gamma}{t_\gamma}(\sum_{\eta=0}^{\alpha-1}\int_{T_\eta}^{T_{\eta+1}} \frac{2\lambda_{M+1,\beta,M}}{\chi_\alpha M} e^{-\int_{T_{\eta+1}}^{T_\alpha} \frac{M\lambda_{M+1,\beta,M}}{\chi_v}dv} e^{-\int_{T_\alpha}^t \frac{M\lambda_{M+1,\beta,M}}{\chi_v}dv} e^{-\int_u^{T_{\eta+1}} \frac{M\lambda_{M+1,\beta,M}}{\chi_v}dv} du + \\
&\qquad \int_{T_\alpha}^t \frac{2\lambda_{M+1,\beta,M}}{\chi_\alpha M} e^{-(t-u)\frac{M\lambda_{M+1,\beta,M}}{\chi_\alpha}dv} du) \\
&= \frac{P_\gamma}{t_\gamma}\frac{2\lambda_{M+1,\beta,M}}{\chi_\alpha M}(\sum_{\eta=0}^{\alpha-1}\int_{T_\eta}^{T_{\eta+1}} e^{-\sum_{\zeta=\eta+1}^{\alpha-1}\Delta_\zeta \frac{M\lambda_{M+1,\beta,M}}{\chi_\zeta}} e^{-(t-T_\alpha)\frac{M\lambda_{M+1,\beta,M}}{\chi_\alpha}} e^{-(T_{\eta+1}-u)\frac{M\lambda_{M+1,\beta,M}}{\chi_\eta}} du \\
&\qquad + \frac{(1-e^{-(t-T_\alpha)\frac{M\lambda_{M+1,\beta,M}}{\chi_\alpha}})}{\frac{M\lambda_{M+1,\beta,M}}{\chi_\alpha}}) \\
&= \frac{P_\gamma}{t_\gamma}\frac{2\lambda_{M+1,\beta,M}}{\chi_\alpha M}(\sum_{\eta=0}^{\alpha-1} e^{-\sum_{\zeta=\eta+1}^{\alpha-1}\Delta_\zeta \frac{M\lambda_{M+1,\beta,M}}{\chi_\zeta}} e^{-(t-T_\alpha)\frac{M\lambda_{M+1,\beta,M}}{\chi_\alpha}} \frac{(1-e^{-\Delta_\eta \frac{M\lambda_{M+1,\beta,M}}{\chi_\eta}})}{\frac{M\lambda_{M+1,\beta,M}}{\chi_\eta}} \\
&\qquad + \frac{(1-e^{-(t-T_\alpha)\frac{M\lambda_{M+1,\beta,M}}{\chi_\alpha}})}{\frac{M\lambda_{M+1,\beta,M}}{\chi_\alpha}})
\end{aligned} \tag{17}
$$

We then have to integrate of the time interval alpha to have the transition probability from the state $\gamma$ to the state $\alpha$.

$$
\begin{aligned}
P(\alpha,i|\gamma,j) &= \int_{T_\alpha}^{T_{\alpha+1}} \frac{P_\gamma}{t_\gamma}\frac{2\lambda_{M+1,\beta,M}}{\chi_\alpha M}(\sum_{\eta=0}^{\alpha-1} e^{-\sum_{\zeta=\eta+1}^{\alpha-1}\Delta_\zeta \frac{M\lambda_{M+1,\beta,M}}{\chi_\zeta}} e^{-(t-T_\alpha)\frac{M\lambda_{M+1,\beta,M}}{\chi_\alpha}} \frac{(1-e^{-\Delta_\eta \frac{M\lambda_{M+1,\beta,M}}{\chi_\eta}})}{\frac{M\lambda_{M+1,\beta,M}}{\chi_\eta}} \\
&\qquad + \frac{(1-e^{-(t-T_\alpha)\frac{M\lambda_{M+1,\beta,M}}{\chi_\alpha}})}{\frac{M\lambda_{M+1,\beta,M}}{\chi_\alpha}})dt \\
&= \frac{P_\gamma}{t_\gamma}\frac{2\lambda_{M+1,\beta,M}}{\chi_\alpha M}(\sum_{\eta=0}^{\alpha-1} e^{-\sum_{\zeta=\eta+1}^{\alpha-1}\Delta_\zeta \frac{M\lambda_{M+1,\beta,M}}{\chi_\zeta}} \frac{(1-e^{-\Delta_\alpha \frac{M\lambda_{M+1,\beta,M}}{\chi_\alpha}})}{\frac{M\lambda_{M+1,\beta,M}}{\chi_\alpha}} \frac{(1-e^{-\Delta_\eta \frac{M\lambda_{M+1,\beta,M}}{\chi_\eta}})}{\frac{M\lambda_{M+1,\beta,M}}{\chi_\eta}} \\
&\qquad + \frac{(\Delta_\alpha - \frac{(1-e^{-(\Delta_\alpha \frac{M\lambda_{M+1,\beta,M}}{\chi_\alpha}})}{\frac{M\lambda_{M+1,\beta,M}}{\chi_\alpha}})}{\frac{M\lambda_{M+1,\beta,M}}{\chi_\alpha}}) \\
&= \frac{P_\gamma}{t_\gamma}\frac{2}{M^2}(\sum_{\eta=0}^{\alpha-1} e^{-\sum_{\zeta=\eta+1}^{\alpha-1}\Delta_\zeta \frac{M\lambda_{M+1,\beta,M}}{\chi_\zeta}} (1-e^{-\Delta_\alpha \frac{M\lambda_{M+1,\beta,M}}{\chi_\alpha}}) \frac{(1-e^{-\Delta_\eta \frac{M\lambda_{M+1,\beta,M}}{\chi_\eta}})}{\frac{M\lambda_{M+1,\beta,M}}{\chi_\eta}} \\
&\qquad + (\Delta_\alpha - \frac{(1-e^{-(\Delta_\alpha \frac{M\lambda_{M+1,\beta,M}}{\chi_\alpha}})}{\frac{M\lambda_{M+1,\beta,M}}{\chi_\alpha}}))
\end{aligned} \tag{18}
$$

Where the recombination probability if defined as:

$$P_\gamma = (1 - e^{-Mrt_\gamma}) \tag{19}$$

$\gamma < \alpha$ We here calculate the transition probability from the state $\gamma$ to a time t in the time interval $\alpha$.

$$\begin{aligned}
P(t,i|t_\gamma,j) = &\int_0^{t_\gamma} \frac{P_\gamma}{t_\gamma} \frac{\lambda_{M,\beta,(M-m+1)}}{\binom{M}{m}\chi_\alpha} e^{-\int_{t_\gamma}^{t} \frac{\lambda_{M,\beta}}{\chi_v} dv} e^{-\int_u^{t_\gamma} \frac{M\lambda_{M+1,\beta,M}}{\chi_v} dv} \frac{2\lambda_{(n+1),\beta,2}}{M(\lambda_{(n+1),\beta,2} + \lambda_{(n+1),\beta,1})} du \\
= &\frac{P_\gamma}{t_\gamma} \frac{\lambda_{M,\beta,(M-m+1)}}{\binom{M}{m}\chi_\alpha} (\sum_{\eta=1}^{\gamma-1} \int_{T_\eta}^{T_{\eta+1}} e^{-\int_{t_\gamma}^{t} \frac{\lambda_{M,\beta}}{\chi_v} dv} e^{-\int_{T_{\eta+1}}^{T_\gamma} \frac{M\lambda_{M+1,\beta,M}}{\chi_v} dv} e^{-\int_{T_\gamma}^{t_\gamma} \frac{M\lambda_{M+1,\beta,M}}{\chi_v} dv} \\
&\times e^{-\int_u^{T_{\eta+1}} \frac{M\lambda_{M+1,\beta,M}}{\chi_v} dv} \frac{2\lambda_{(n+1),\beta,2}}{M(\lambda_{(n+1),\beta,2} + \lambda_{(n+1),\beta,1})} du \\
&+ \int_{T_\gamma}^{t_\gamma} e^{-\int_{t_\gamma}^{t} \frac{\lambda_{M,\beta}}{\chi_v} dv} e^{-\int_u^{t_\gamma} \frac{M\lambda_{M+1,\beta,M}}{\chi_v} dv} \frac{2\lambda_{(n+1),\beta,2}}{M(\lambda_{(n+1),\beta,2} + \lambda_{(n+1),\beta,1})} du) \\
= &\frac{P_\gamma}{t_\gamma} \frac{2\lambda_{(n+1),\beta,2} e^{-\int_{t_\gamma}^{t} \frac{\lambda_{M,\beta}}{\chi_v} dv}}{M(\lambda_{(n+1),\beta,2} + \lambda_{(n+1),\beta,1})} \frac{\lambda_{M,\beta,(M-m+1)}}{\binom{M}{m}\chi_\alpha} (\sum_{\eta=1}^{\gamma-1} e^{-\sum_{\zeta=\eta+1}^{\gamma-1} \Delta_\zeta \frac{M\lambda_{M+1,\beta,M}}{\chi_\zeta}} \\
&\times e^{-(t_\gamma - T_\gamma)\frac{M\lambda_{M+1,\beta,M}}{\chi_\gamma}} \frac{(1 - e^{-\Delta_\eta \frac{M\lambda_{M+1,\beta,M}}{\chi_\eta}})}{\frac{M\lambda_{M+1,\beta,M}}{\chi_\eta}} \\
&+ \frac{(1 - e^{-(t_\gamma - T_\gamma)\frac{M\lambda_{M+1,\beta,M}}{\chi_\gamma}})}{\frac{M\lambda_{M+1,\beta,M}}{\chi_\gamma}})
\end{aligned} \tag{20}$$

We then have to integrate of the time interval alpha to have the transition probability from the state $\gamma$ to the state $\alpha$.

$$\begin{aligned}
P(\alpha,i|\gamma,j) = &\int_{T_\alpha}^{T_{\alpha+1}} \frac{P_\gamma}{t_\gamma} \frac{2\lambda_{(n+1),\beta,2} e^{-\int_{t_\gamma}^{t} \frac{\lambda_{M,\beta}}{\chi_v} dv}}{M(\lambda_{(n+1),\beta,2} + \lambda_{(n+1),\beta,1})} \frac{\lambda_{M,\beta,(M-m+1)}}{\binom{M}{m}\chi_\alpha} \\
&(\sum_{\eta=1}^{\gamma-1} e^{-\sum_{\zeta=\eta+1}^{\gamma-1} \Delta_\zeta \frac{M\lambda_{M+1,\beta,M}}{\chi_\zeta}} e^{-(t_\gamma - T_\gamma)\frac{M\lambda_{M+1,\beta,M}}{\chi_\gamma}} \frac{(1 - e^{-\Delta_\eta \frac{M\lambda_{M+1,\beta,M}}{\chi_\eta}})}{\frac{M\lambda_{M+1,\beta,M}}{\chi_\eta}} + \frac{(1 - e^{-(t_\gamma - T_\gamma)\frac{M\lambda_{M+1,\beta,M}}{\chi_\gamma}})}{\frac{M\lambda_{M+1,\beta,M}}{\chi_\gamma}})dt \\
= &\int_{T_\alpha}^{T_{\alpha+1}} \frac{P_\gamma}{t_\gamma} \frac{2\lambda_{(n+1),\beta,2} e^{-\int_{t_\gamma}^{T_\alpha} \frac{\lambda_{M,\beta}}{\chi_v} dv} e^{-\int_{T_\alpha}^{t} \frac{\lambda_{M,\beta}}{\chi_v} dv}}{M(\lambda_{(n+1),\beta,2} + \lambda_{(n+1),\beta,1})} \frac{\lambda_{M,\beta,(M-m+1)}}{\binom{M}{m}\chi_\alpha} \\
&(\sum_{\eta=1}^{\gamma-1} e^{-\sum_{\zeta=\eta+1}^{\gamma-1} \Delta_\zeta \frac{M\lambda_{M+1,\beta,M}}{\chi_\zeta}} e^{-(t_\gamma - T_\gamma)\frac{M\lambda_{M+1,\beta,M}}{\chi_\gamma}} \frac{(1 - e^{-\Delta_\eta \frac{M\lambda_{M+1,\beta,M}}{\chi_\eta}})}{\frac{M\lambda_{M+1,\beta,M}}{\chi_\eta}} + \frac{(1 - e^{-(t_\gamma - T_\gamma)\frac{M\lambda_{M+1,\beta,M}}{\chi_\gamma}})}{\frac{M\lambda_{M+1,\beta,M}}{\chi_\gamma}})dt \\
= &\frac{P_\gamma}{t_\gamma} \frac{2\lambda_{(n+1),\beta,2} e^{-\int_{t_\gamma}^{T_\alpha} \frac{\lambda_{M,\beta}}{\chi_v} dv} (1 - e^{-\Delta_\alpha \frac{\lambda_{M,\beta}}{\chi_\alpha}})}{M(\lambda_{(n+1),\beta,2} + \lambda_{(n+1),\beta,1})} \frac{\lambda_{M,\beta,(M-m+1)}}{\binom{M}{m}\lambda_{M,\beta}} \\
&(\sum_{\eta=1}^{\gamma-1} e^{-\sum_{\zeta=\eta+1}^{\gamma-1} \Delta_\zeta \frac{M\lambda_{M+1,\beta,M}}{\chi_\zeta}} e^{-(t_\gamma - T_\gamma)\frac{M\lambda_{M+1,\beta,M}}{\chi_\gamma}} \frac{(1 - e^{-\Delta_\eta \frac{M\lambda_{M+1,\beta,M}}{\chi_\eta}})}{\frac{M\lambda_{M+1,\beta,M}}{\chi_\eta}} + \frac{(1 - e^{-(t_\gamma - T_\gamma)\frac{M\lambda_{M+1,\beta,M}}{\chi_\gamma}})}{\frac{M\lambda_{M+1,\beta,M}}{\chi_\gamma}})
\end{aligned} \tag{21}$$

$\gamma = \alpha, m = n + 1$

For a multiple merger event to happen, there are three possibilities. A non coalescing branch join the coalescent event, or it coalesces in the same hidden state (before or after the coalescent event).

$$
\begin{aligned}
P(\gamma, i|\gamma, j) =& \frac{P_\gamma}{t_\gamma}\int_0^{t_\gamma} \frac{(M-n)\lambda_{(n+1),\beta,1} e^{-\int_u^{t_\gamma} \frac{M\lambda_{M+1,\beta,M}}{\chi v} dv}}{M(\lambda_{(n+1),\beta,2}+\lambda_{(n+1),\beta,1})} du + Pc_1 + Pc_2 \\
=& \frac{P_\gamma}{t_\gamma}\frac{(M-n)\lambda_{(n+1),\beta,1}}{M(\lambda_{(n+1),\beta,2}+\lambda_{(n+1),\beta,1})}(\sum_{\eta=1}^{\gamma-1}\int_{T_\eta}^{T_{\eta+1}} e^{-\int_u^{T_{\eta+1}} \frac{M\lambda_{M+1,\beta,M}}{\chi v} dv} e^{-\int_{T_{\eta+1}}^{t_\gamma} \frac{M\lambda_{M+1,\beta,M}}{\chi v} dv} du \\
&+ \int_{T_\gamma}^{t_\gamma} e^{-\int_u^{t_\gamma} \frac{M\lambda_{M+1,\beta,M}}{\chi v} dv} du) + Pc_1 + Pc_2 \\
=& \frac{P_\gamma}{t_\gamma}\frac{(M-n)\lambda_{(n+1),\beta,1}}{M(\lambda_{(n+1),\beta,2}+\lambda_{(n+1),\beta,1})}(\sum_{\eta=1}^{\gamma-1}\frac{(1-e^{-\Delta_\eta \frac{M\lambda_{M+1,\beta,M}}{\chi\eta}})}{\frac{M\lambda_{M+1,\beta,M}}{\chi\eta}} e^{-\int_{T_{\eta+1}}^{t_\gamma} \frac{M\lambda_{M+1,\beta,M}}{\chi v} dv} \\
&+ \frac{(1-e^{-(t_\gamma - T_\gamma)\frac{M\lambda_{M+1,\beta,M}}{\chi\gamma}})}{\frac{M\lambda_{M+1,\beta,M}}{\chi\gamma}}) + Pc_1 + Pc_2 \\
=& \frac{P_\gamma(M-n)\lambda_{(n+1),\beta,1}}{t_\gamma M(\lambda_{(n+1),\beta,2}+\lambda_{(n+1),\beta,1})}(\sum_{\eta=1}^{\gamma-1}\frac{(1-e^{-\Delta_\eta \frac{M\lambda_{M+1,\beta,M}}{\chi\eta}})}{\frac{M\lambda_{M+1,\beta,M}}{\chi\eta}} e^{-\sum_{\zeta=\eta+1}^{\gamma-1}\frac{\Delta_\zeta M\lambda_{M+1,\beta,M}}{\chi\zeta}} e^{-(t_\gamma - T_\gamma)\frac{M\lambda_{M+1,\beta,M}}{\chi\gamma}} \\
&+ \frac{(1-e^{-(t_\gamma - T_\gamma)\frac{M\lambda_{M+1,\beta,M}}{\chi\gamma}})}{\frac{M\lambda_{M+1,\beta,M}}{\chi\gamma}}) + Pc_1 + Pc_2 \qquad (22)
\end{aligned}
$$

$Pc_1$ is the probability that a recombination happens before the first coalescent event in the non coalescing branch, it then coalesce before the current first coalescent event but in the same hidden state (resulting in a multiple merger coalescent because of the discretized time)

$Pc_2$ is the probability that a recombination happens before $T_{\gamma+1}$ in the non coalescing branch, it then coalesce after the current first coalescent event but in the same hidden state (resulting in a multiple merger coalescent because of the discretized time)

$$Pc_1 = \int_{T_\gamma}^{t_\gamma} \frac{P_\gamma}{t_\gamma}\frac{\lambda_{2,\beta}}{\chi_\gamma M}(\sum_{\eta=0}^{\gamma-1} e^{-\sum_{\zeta=\eta+1}^{\gamma-1}\Delta_\zeta \frac{M\lambda_{M+1,\beta,M}}{\chi_\zeta}} e^{-(t-T_\gamma)\frac{M\lambda_{M+1,\beta,M}}{\chi_\gamma}} \frac{(1-e^{-\Delta_\eta \frac{M\lambda_{M+1,\beta,M}}{\chi_\eta}})}{\frac{M\lambda_{M+1,\beta,M}}{\chi_\eta}} + \frac{(1-e^{-(t-T_\gamma)\frac{M\lambda_{M+1,\beta,M}}{\chi_\gamma}})}{\frac{M\lambda_{M+1,\beta,M}}{\chi_\gamma}})dt$$

$$= \frac{P_\gamma}{t_\gamma}\frac{\lambda_{2,\beta}}{\chi_\gamma M}(\sum_{\eta=0}^{\gamma-1} e^{-\sum_{\zeta=\eta+1}^{\gamma-1}\Delta_\zeta \frac{M\lambda_{M+1,\beta,M}}{\chi_\zeta}} \frac{(1-e^{-(t_\gamma-T_\gamma)\frac{M\lambda_{M+1,\beta,M}}{\chi_\gamma}})}{\frac{M\lambda_{M+1,\beta,M}}{\chi_\gamma}} \frac{(1-e^{-\Delta_\eta \frac{M\lambda_{M+1,\beta,M}}{\chi_\eta}})}{\frac{M\lambda_{M+1,\beta,M}}{\chi_\eta}} + \frac{((t_\gamma-T_\gamma) - \frac{(1-e^{-(t_\gamma-T_\gamma)\frac{M\lambda_{M+1,\beta,M}}{\chi_\gamma}})}{\frac{M\lambda_{M+1,\beta,M}}{\chi_\gamma}})}{\frac{M\lambda_{M+1,\beta,M}}{\chi_\gamma}}) \tag{23}$$

$$= \frac{P_\gamma}{t_\gamma}\frac{1}{M^2}(\sum_{\eta=0}^{\gamma-1} e^{-\sum_{\zeta=\eta+1}^{\gamma-1}\Delta_\zeta \frac{M\lambda_{M+1,\beta,M}}{\chi_\zeta}} (1-e^{-(t_\gamma-T_\gamma)\frac{M\lambda_{M+1,\beta,M}}{\chi_\gamma}}) \frac{(1-e^{-\Delta_\eta \frac{M\lambda_{M+1,\beta,M}}{\chi_\eta}})}{\frac{M\lambda_{M+1,\beta,M}}{\chi_\eta}} +((t_\gamma-T_\gamma) - \frac{(1-e^{-(t_\gamma-T_\gamma)\frac{M\lambda_{M+1,\beta,M}}{\chi_\gamma}})}{\frac{M\lambda_{M+1,\beta,M}}{\chi_\gamma}}))$$

$$Pc_2 = \int_{t_\gamma}^{T_{\gamma+1}} \frac{P_\gamma}{t_\gamma}\frac{\lambda_{(n+1),\beta,2} e^{-\int_{t_\gamma}^{t}\frac{\lambda_{2,\beta}}{\chi_\gamma}dv}}{M(\lambda_{(n+1),\beta,2}+\lambda_{(n+1),\beta,1})}\frac{\lambda_{2,\beta}}{\chi_\gamma} (\sum_{\eta=1}^{\gamma-1} e^{-\sum_{\zeta=\eta+1}^{\gamma-1}\Delta_\zeta \frac{M\lambda_{M+1,\beta,M}}{\chi_\zeta}} e^{-(t_\gamma-T_\gamma)\frac{M\lambda_{M+1,\beta,M}}{\chi_\gamma}} \frac{(1-e^{-\Delta_\eta \frac{M\lambda_{M+1,\beta,M}}{\chi_\eta}})}{\frac{M\lambda_{M+1,\beta,M}}{\chi_\eta}} + \frac{(1-e^{-\Delta_\gamma \frac{M\lambda_{M+1,\beta,M}}{\chi_\gamma}})}{\frac{M\lambda_{M+1,\beta,M}}{\chi_\gamma}})dt$$

$$= \frac{P_\gamma}{t_\gamma}\frac{\lambda_{(n+1),\beta,2}(1-e^{-(T_{\gamma+1}-t_\gamma)\frac{\lambda_{2,\beta}}{\chi_\gamma}})}{M(\lambda_{(n+1),\beta,2}+\lambda_{(n+1),\beta,1})} (\sum_{\eta=1}^{\gamma-1} e^{-\sum_{\zeta=\eta+1}^{\gamma-1}\Delta_\zeta \frac{M\lambda_{M+1,\beta,M}}{\chi_\zeta}} e^{-(t_\gamma-T_\gamma)\frac{M\lambda_{M+1,\beta,M}}{\chi_\gamma}} \frac{(1-e^{-\Delta_\eta \frac{M\lambda_{M+1,\beta,M}}{\chi_\eta}})}{\frac{M\lambda_{M+1,\beta,M}}{\chi_\eta}} + \frac{(1-e^{-\Delta_\gamma \frac{M\lambda_{M+1,\beta,M}}{\chi_\gamma}})}{\frac{M\lambda_{M+1,\beta,M}}{\chi_\gamma}}) \tag{24}$$

$\gamma = \alpha, m = n-1$

$$
\begin{aligned}
P(\gamma,i|\gamma,j) &= \frac{P_\gamma}{t_\gamma}\int_0^{t_\gamma}\frac{\lambda_{(n+1),\beta,2}e^{-\int_u^{t_\gamma}\frac{M\lambda_{M+1,\beta,M}}{\chi v}dv}}{n(\lambda_{(n+1),\beta,2}+\lambda_{(n+1),\beta,1})}du \\
&= \frac{P_\gamma}{t_\gamma}\frac{\lambda_{(n+1),\beta,2}}{n(\lambda_{(n+1),\beta,2}+\lambda_{(n+1),\beta,1})}\Big(\sum_{\eta=1}^{\gamma-1}\int_{T_\eta}^{T_{\eta+1}} e^{-\int_u^{T_{\eta+1}}\frac{M\lambda_{M+1,\beta,M}}{\chi v}dv}e^{-\int_{T_{\eta+1}}^{t_\gamma}\frac{M\lambda_{M+1,\beta,M}}{\chi v}dv}du \\
&\quad + \int_{T_\gamma}^{t_\gamma} e^{-\int_u^{t_\gamma}\frac{M\lambda_{M+1,\beta,M}}{\chi v}dv}du\Big) \\
&= \frac{P_\gamma}{t_\gamma}\frac{\lambda_{(n+1),\beta,2}}{n(\lambda_{(n+1),\beta,2}+\lambda_{(n+1),\beta,1})}\Big(\sum_{\eta=1}^{\gamma-1}\frac{(1-e^{-\Delta_\eta\frac{M\lambda_{M+1,\beta,M}}{\chi\eta}})}{\frac{M\lambda_{M+1,\beta,M}}{\chi\eta}}e^{-\int_{T_{\eta+1}}^{t_\gamma}\frac{M\lambda_{M+1,\beta,M}}{\chi v}dv}+\frac{(1-e^{-(t_\gamma-T_\gamma)\frac{M\lambda_{M+1,\beta,M}}{\chi\gamma}})}{\frac{M\lambda_{M+1,\beta,M}}{\chi\gamma}}\Big) \\
&= \frac{P_\gamma}{t_\gamma}\frac{\lambda_{(n+1),\beta,2}}{n(\lambda_{(n+1),\beta,2}+\lambda_{(n+1),\beta,1})}\Big(\sum_{\eta=1}^{\gamma-1}\frac{(1-e^{-\Delta_\eta\frac{M\lambda_{M+1,\beta,M}}{\chi\eta}})}{\frac{M\lambda_{M+1,\beta,M}}{\chi\eta}}e^{-\sum_{\zeta=\eta+1}^{\gamma-1}\Delta_\zeta\frac{M\lambda_{M+1,\beta,M}}{\chi\zeta}}e^{-(t_\gamma-T_\gamma)\frac{M\lambda_{M+1,\beta,M}}{\chi\gamma}} \\
&\quad +\frac{(1-e^{-(t_\gamma-T_\gamma)\frac{M\lambda_{M+1,\beta,M}}{\chi\gamma}})}{\frac{M\lambda_{M+1,\beta,M}}{\chi\gamma}}\Big)
\end{aligned}
\tag{25}
$$

$\gamma = \alpha, m = n$

$$P(\gamma,j|\gamma,j) = 1-\sum_{\alpha\neq\gamma,i\neq j}P(\alpha,i|\gamma,j) \tag{26}$$

#### 1.5 Emission Matrix

To compute the emission probabilities, we need the external branch length at each position of the genome (noted as $Ts$). $Ts$ is estimated with an additional HMM as described in [4].

### 1.5.1 M=3

One can only observe 4 possibilities. Observation 0, no mutation within the 3 individual. Observation 1 (2,3), individual 1 (2,3) is different from individual 2 and 3 (1 and 3 , 1 and 2).

if the first coalescent event involves 2 individuals we have :

$$P(0|\gamma) = e^{-\mu(Ts)} \tag{27}$$

$i \in 1,2,3$. Mutation occurred and did not occurred in the first coalescent event.

$$P(i|\gamma,i) = (1-e^{-\mu(Ts-2t_\gamma)}) \tag{28}$$

$i \in 1,2,3; \bar{i} \neq i$. Mutation occurred and is in the first coalescent event.

$$P(i|\gamma,\bar{i}) = (1-e^{-\mu(2t_\gamma)}) \tag{29}$$

if the first coalescent event involves 3 individuals we have :

$$P(0|\gamma) = e^{-\mu(3t_\gamma)} \tag{30}$$

$i \in 1,2,3$

$$P(i|\gamma) = 1-e^{-\mu(3t_\gamma)} \tag{31}$$

### 1.5.2 M=4

One can only observe 8 possibilities as we only focus on SNPs. Observation 0, no mutation within the 4 individual. Observation 1 (2,3,4), individual 1 (2,3,4) is different from all other individual. Observation 5 to 7 , two individual are different from the other two.

if the first coalescent event involves 4 individuals:

$$P(0|\gamma) = e^{-\mu(4t_\gamma)} \tag{32}$$

$i \in 1, 2, 3, 4$

$$P(i|\gamma) = 1 - e^{-\mu(4t_\gamma)} \tag{33}$$

if the first coalescent event involves 3 individuals:

$$P(0|\gamma) = e^{-\mu(Ts)} \tag{34}$$

$i \in 1, 2, 3, 4$

$$P(i|\gamma, \bar{i}) = (1 - e^{-\mu(t_\gamma)}) \tag{35}$$

$i \in 1, 2, 3, 4$

$$P(i|\gamma, i) = (1 - e^{-\mu(Ts-3t_\gamma)}) \tag{36}$$

if the first coalescent event involves 2 individuals:
No mutation occurred.

$$P(0|\gamma) = e^{-\mu(Ts)} \tag{37}$$

$i \in 1, 2, 3, 4$ and mutation is within on one of the individual that coalesce.

$$P(i|\gamma) = (1 - e^{-\mu(t_\gamma)}) \tag{38}$$

$i \in 1, 2, 3, 4$ and mutation is not on one of the individual that coalesce.

$$P(i|\gamma) = (1 - e^{-\mu(\frac{(Ts-4t_\gamma)}{2}+t_\gamma)}) \tag{39}$$

$i \in 5, 6, 7$ and the two individual coalescing are identical.

$$P(i|\gamma) = e^{-\mu(Ts)} \tag{40}$$

$i \in 5, 6, 7$ and the two individual coalescing are different,then two mutation must occur, which has probability 0.

$$P(i|\gamma) = 0 \tag{41}$$
