## Supplementary Text S2 for "Simultaneous Inference of Past Demography and Selection from the Ancestral Recombination Graph under the Beta Coalescent"

### **Supplementary Text S2:** Description of the Graph Neural Network Approach (GNN*coal*)

### 1 Brief Introduction to neural network

We wish to define a neural network that has some learnable parameters (i.e. weights) to predict parameters of interest. In this study, we focus on the past variation of population size and on the $\alpha$ parameter of the Beta distribution shaping the offspring distribution [8]. To define a neural network, we first generate a training dataset, *i.e.* a dataset containing information to learn from (the ancestral recombination graph) and predictive variables of interest (past demography and $\alpha$ parameter). We then process inputs through the neural network and compute the loss, which evaluates the distance between predictions and ground truth. Afterwards, we propagate gradients back into the network's parameters and update its weights. We now describe further details of this procedure.

### 2 Datasets

#### 2.1 Set 1: Variable demography and MMC

To build our training data set, we rely on the efficient coalescent and sequence simulator *msprime* [1, 4]. We simulated 10 haploid genomes for a total of $2 \times 10^4$ demographic scenarios with 100 replicates each. We followed the procedure outlined in Figure S1 in Supplementary text S2 (see below) to define our scenarios. To ensure that each simulation contains sufficient data points, we imposed that 95% of the 100 replicates had to contain at least 500 trees. This procedure resulted in a training data set containing between $9.5 \times 10^8$ and $1 \times 10^9$ coalescent trees. Each simulated data point was stored as a tree sequence and converted into the graph format (*i.e. Pytorch Geometric* data object for the forward pass. This procedure drastically decreased the storage space requirements at the marginal cost of training speed. Furthermore, we trained our network over a range of the multiple merger parameter $\alpha$ from 1.01 to 1.99.

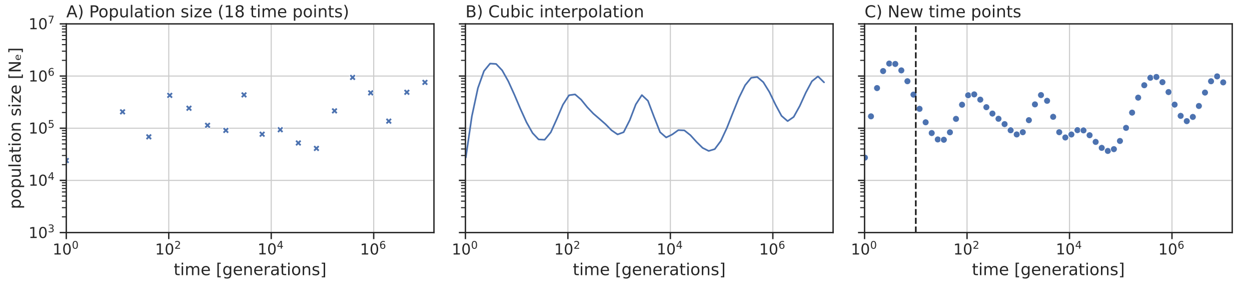


**Fig. S1. Demography sampling process** Population size (y-axis) changes through time in generations (x-axis). Demography sampling process according to [2, 7] (A) and cubic interpolation (B), and re-sampling for 60 uniformly spaced points. The sampling process is repeated if any population size is outside the $10^4$ and $10^7$ population size window.

#### 2.2 Set 2: MMC versus selection under constant population size

To discern between multiple merger and selection, we simulated 1000 replicates under constant population size ($N_e = 10^5$) by randomly choosing between Kingman and $\beta$-coalescent models. Additionally, we randomly chose one of the selection regimes when a

Kingman coalescent scenario was drawn. For each scenario, we simulated 100 replicates with 99% of each replicate containing at least 2,000 trees (*i.e.* a batch size of 2000). This procedure resulted in a training data set of more than $1.98 \times 10^8$ coalescent trees. An additional validation dataset was generated consisting of 1,000 simulations under the same scenarios. A script for replicating these simulations is available at `https://github.com/kevinkorfmann/GNNcoal-analysis`.

### 3 Training

The demography inference model was trained for two epochs, an epoch being defined by iterating one time over the training dataset. *i.e.* the data is processed twice, once by the untrained neural network and a second time after the first epoch. The model to infer the $\alpha$ parameter was trained on 10% of the dataset used for demography inference as we found the procedure to be sufficient for accurate estimates. We set the batch size to 500 (*i.e.* number of trees processed before updating the neural network parameters). Jupyter notebooks to perform the training are accessible at `https://github.com/kevinkorfmann/GNNcoal-analysis`.

### 4 Neural network

The neural network is implemented in *PyTorch* [6] using the extension *Pytorch Geometric* for the graph convolution operation and differential hierarchical pooling strategy [3, 5, 9]. For training and inference, each coalescent tree was interpreted as an undirected graph, with each ancestral node or leaf having edges to their parent or children, and vice versa. Each node $i$ contains a feature vector (a feature vector being the predicted values by the neural network) $\mathbf{x_i}$ of size 60 (number of desired estimated parameters), which are horizontally stacked into a feature vector matrix (*i.e.* each node/coalescent event has feature vector) $\mathbf{X}$ and initialized with ones on the diagonal and zeros everywhere else. Each feature vector are unique. As mentioned above, we chose a feature vector of size 60, which corresponds to the final output dimension of the demography network. A convolution of an individual feature vector $\mathbf{x_i}$ of node $i$ is computed following [5]:

$$\mathbf{x}'_i = \mathbf{\Theta}^\top \sum_{j \in \mathcal{N}(v) \cup \{i\}} \frac{e_{j,i}}{\sqrt{\hat{d}_j \hat{d}_i}} \mathbf{x}_j \tag{1}$$

With $\hat{d}_i = 1 + \sum_{j \in \mathcal{N}(i)} e_{j,i}$ and $e_{j,i}$ being the pivot edge from $j$ to $i$. The node features $\mathbf{x_i}$ are transformed by a learnable weight matrix $\Theta$, normalized according to the degrees of the respective nodes, and summed over the neighboring nodes. Alongside this node-wise formulation, the same computation can be written as matrix-operations. The latter will be the notation used to explain the following pooling strategy:

$$\mathbf{X}' = \hat{\mathbf{D}}^{-1/2} \hat{\mathbf{A}} \hat{\mathbf{D}}^{-1/2} \mathbf{X} \mathbf{\Theta}, \tag{2}$$

Where $\hat{\mathbf{A}} = \mathbf{A} + \mathbf{I}$ represents the adjacency matrix and $\hat{D}_{ii} = \sum_{j=0} \hat{A}_{ij}$, represent the diagonal degree matrix. Here, the edge information enters the convolution as part of the values of the adjacency matrix. Both representations are valid descriptions of the same convolution [3].

We defined our GNN as a series of three consecutive graph convolutions, with input, hidden and output dimensions of size 60, each convolution separated by a one-dimensional batch normalization step. Increasing the number of convolutions above a certain value will lead to a *smoothing* of the feature vectors, as the information of more distant neighbours will be taken into account for each node [5]. The output of each convolution is then concatenated and passed through-linear activation and ReLu-function ($Relu(z) = max(0, z)$). The complete implementation is available at `https://github.com/kevinkorfmann/GNNcoal`.

Because each node has a feature vector (*i.e.* each coalescent event of a coalescent tree lead to a specific prediction), the feature vector matrix (*i.e.* the collection of estimated demography parameters from a coalescent tree) needs to be processed to obtain a consensus predicted demography. This step is called the pooling strategy. A detailed description of the pooling strategy used in our study is formulated in [9] and short description can be found below.

We compress the number of nodes hierarchically in a step-wise manner, from the $l$ state to $l+1$ state. Close nodes in the original graph are summarized by a supernode in a new smaller graph, ideally having only one supernode at the end with one feature vector, containing our variables of interest. Thus, this downsizing can be formulated as decreasing the original adjacency matrix $\mathbf{A}$ of a given genealogy to a smaller subgraph $\mathbf{A}^{l+1}$ ($\mathbf{A} \in \mathbb{R}^{n \times n} \rightarrow \mathbf{A}^{l+1} \in \mathbb{R}^{m \times m}$ with $n, m$ being the number of nodes in each graph and $\mathbf{Z} \in \mathbb{R}^{n \times d} \rightarrow \mathbf{Z}^{l+1} \in \mathbb{R}^{m \times d}$ with condition of $m < n$; here $d$ being the length of the feature vector). To achieve this, we cluster a set of nodes in the original genealogy into a smaller subset of supernodes. A cluster assignment matrix $\mathbf{S} \in \mathbb{R}^{n^l \times n^{l+1}}$ (with $n^l$ is the number of clusters in the current step and $n^{l+1}$ the number of clusters in the next step). In the first iteration, the number of clusters $n^l$ is equal to the number of nodes, and it is *learned* using an $GNN_{pool}$. This means that the criterion to compress individual nodes is dependent on the overall objective of network, which is to reduce the loss function with respect to predicting parameters of interest. The specific computation using two GNNs (($GNN_{pool}$ and $GNN_{embed}$)) is explained in the following:

1. We compute the feature vectors $\mathbf{Z^l}$ using $GNN_{embed}$ at step $l$ (equation (3)).

$$\mathbf{Z}^l = GNN_{l,embed}(\mathbf{A}^l, \mathbf{X}^l) \tag{3}$$

2. Next we compute the cluster assignment matrix, by computing "feature vectors" and applying the row-wise softmax function ($\sigma(z_i) = \frac{e^{z_i}}{\sum_{j=1}^{d} e^{z_j}} \quad for \ i = 1, 2, \dots, d$) as normalization step. This results is a probability distribution of each node to be part of the next cluster.

$$\mathbf{S}^l = softmax(GNN_{l,pool}(\mathbf{A}^l, \mathbf{X}^l)) \tag{4}$$

3. Lastly, the new adjacency matrix $\mathbf{A}^{l+1}$ and corresponding feature vector matrix $\mathbf{X}^{l+1}$ are calculated ( (5) and (6)). Notably, in equation $\mathbf{A^l}$ is multiplied twice with $\mathbf{S}$ to ensure that $\mathbf{A}^{l+1}$ is square (*i.e.* the number of columns is equal to the number of rows).

$$\mathbf{X}^{l+1} = \mathbf{S}^{l^T} \mathbf{Z}^l \rightarrow \mathbb{R}^{n^{l+1} \times d} \tag{5}$$

$$\mathbf{A}^{l+1} = \mathbf{S}^{l^T} \mathbf{A^l} \mathbf{S}^l \rightarrow \mathbb{R}^{n^{l+1} \times n^{l+1}} \tag{6}$$

In our approach, we used three consecutive pooling iterations $l = 1, ..., 3$. At each iteration, a graph is compressed down to 30% of the remaining nodes. In total, six GNNs (three embedding GNNs and three pooling GNNs) contribute to the number of learnable parameters, followed by two linear layers with interspaced ReLU-activation to compress the output into the desired demographic-time window dimension. The demography model implementation can be found at `https://github.com/kevinkorfmann/GNNcoal`.

We now focus on the $\alpha$ inference and the classification. The model for inferring the $\alpha$ parameter adds three linear layers to the demography network model, while the classification model adds two linear layers. To illustrate the simplicity to estimate $\alpha$ and classify models, we show the code for reducing the number of dimensions below.

$\alpha$ inference model:

```
class AlphaInferenceModel(nn.Module):

    def __init__(self, DemographyNet, time_window=60):
        super().__init__()
        self.l1 = nn.Linear(time_window, time_window//2)
        self.l2 = nn.Linear(time_window//2, time_window//4)
        self.l3 = nn.Linear(time_window//4, 1)
        self.DemographyNet = DemographyNet

    def forward(self, batch):
        x = self.DemographyNet(batch)
        return self.l3(F.relu(self.l2(F.relu(self.l1(x)))))
```

Classification model:

```
class ClassificationModel(nn.Module):

    def __init__(self, DemographyNet, num_classes, time_window=60):
        super().__init__()
        self.l1 = nn.Linear(time_window, time_window//2)
        self.l2 = nn.Linear(time_window//2, num_classes)
        self.DemographyNet = DemographyNet

    def forward(self, batch):
        x = self.DemographyNet(batch)
        return self.l2(F.relu(self.l1(x)))
```

### 5 Time window

The output of the neural network is a fixed-size vector containing the values of inferred population size. However, not all trees contain coalescent events falling into these pre-specified time-bins. As a consequence, the population size can only be inferred for simulations where trees contain coalescence events in those specific bins. A multi-step heuristic has been developed to obtain an estimate of the inference window for each simulation, respectively. The procedure is summarized as:

```
1. Simulate 100 repetitions for a demographic parameter set.
2. Count number of coalescent events for first 500 trees for each simulation.
3. Smoothen coalescent event vectors with sliding window*.
4. Only retain the largest consecutive coalescent time-window per repetition.
5. Create Boolean mask by retaining the time-windows with at least X coalescent events
along repetition axis.

*Sliding window: If left and right of pivot element up to Y coalescent events
are present, retain the pivot element, move one step to the right and repeat.
```

This heuristically constructed mask is used to zero out all inferences outside of the coalescent events before computing the RMSE-loss function and back-propagating the weights (avoiding under- and over-fitting). In practice, a single row of the mask, is multiplied into a matrix of dimensions equal to the number of trees times the number of windows (here: 500x60). With this procedure, each tree receives the same masked loss based on all repetitions (to balance the training data set). Coalescent events and masks for constant demography of size $10^6$ are provided in Figure S2 below.

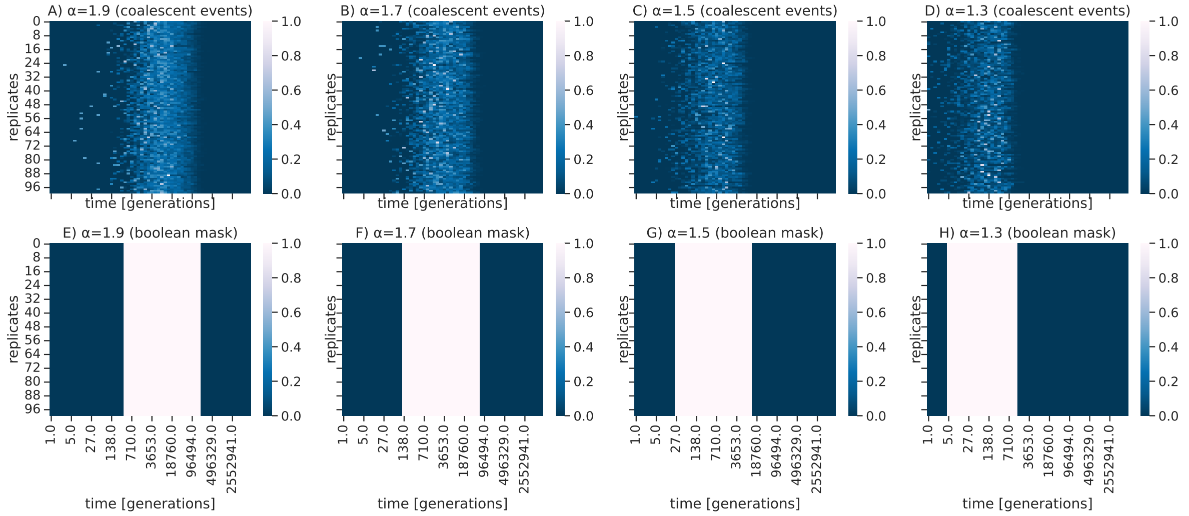


**Fig. S2. Coalescent events and boolean masks** A)-D) The number of coalescent events for 100 repetitions (y-axis) across the first 500 trees of a simulation for 60 time windows (x-axis). E)-H): Boolean mask after applying a heuristic to determine a suitable time-window for demography inference.

During the inference, when only one sample is available, the masking procedure was approximated by calculating the mean of the log-scaled node times of the first 500 trees and by taking two standard deviations at both sides from the mean to form a similar time-window even if only one repetition is available (see below).

```
def alternative_coalescent_mask(ts, population_time, x_times_std=2, n_trees=500):

    trees = ts.aslist()[0:n_trees]
    nodes_n_trees = []
    for tree in trees:
        nodes_n_trees += list(tree.nodes())

    node_times = [ts.get_time(node.id) for node in ts.nodes() if node.id >= ts.num_samples and
        node.id in nodes_n_trees]

    log_node_times = np.log(node_times)
    mean = log_node_times.mean()
    std = log_node_times.std()
    lowerbound = np.exp(mean-x_times_std*std)
    upperbound = np.exp(mean+x_times_std*std)
    mask1 = population_time > lowerbound
    mask2 = population_time < upperbound
    mask = np.logical_and(mask1, mask2)
    return mask
```

### References

[1] Franz Baumdicker, Gertjan Bisschop, Daniel Goldstein, Graham Gower, Aaron P. Ragsdale, Georgia Tsambos, Sha Zhu, Bjarki Eldon, E. Castedo Ellerman, Jared G. Galloway, Ariella L. Gladstein, Gregor Gorjanc, Bing Guo, Ben Jeffery, Warren W. Kretzschumar, Konrad Lohse, Michael Matschiner, Dominic Nelson, Nathaniel S. Pope, Consuelo D. Quinto-Cortes, Murillo F. Rodrigues, Kumar Saunack, Thibaut Sellinger, Kevin Thornton, Hugo van Kemenade, Anthony W. Wohns, Yan Wong, Simon Gravel, Andrew D. Kern, Jere Koskela, Peter L. Ralph, and Jerome Kelleher. Efficient ancestry and mutation simulation with msprime 1.0. *GENETICS*, 220(3), MAR 3 2022.

[2] Simon Boitard, Willy Rodríguez, Flora Jay, Stefano Mona, and Frédéric Austerlitz. Inferring population size history from large samples of genome-wide molecular data - an approximate bayesian computation approach. 12(3):e1005877.

[3] Matthias Fey and Jan Eric Lenssen. Fast graph representation learning with PyTorch geometric.

[4] Jerome Kelleher, Kevin R. Thornton, Jaime Ashander, and Peter L. Ralph. Efficient pedigree recording for fast population genetics simulation. 14(11):e1006581.

[5] Thomas N. Kipf and Max Welling. Semi-Supervised Classification with Graph Convolutional Networks. 2016.

[6] Adam Paszke, Sam Gross, Soumith Chintala, Gregory Chanan, Edward Yang, Zachary DeVito, Zeming Lin, Alban Desmaison, Luca Antiga, and Adam Lerer. Automatic differentiation in PyTorch. October 2017.

[7] Théophile Sanchez, Jean Cury, Guillaume Charpiat, and Flora Jay. Deep learning for population size history inference: Design, comparison and combination with approximate bayesian computation. 21(8):2645–2660.

[8] J Schweinsberg. Coalescent processes obtained from supercritical Galton-Watson processes. *Stochastic Processes and their Applications*, 106(1):107–139, JUL 2003.

[9] Rex Ying, Jiaxuan You, Christopher Morris, Xiang Ren, William L. Hamilton, and Jure Leskovec. Hierarchical graph representation learning with differentiable pooling.
